# Robust Prediction of Multiple Protein Conformations with Entropy Guidance

**DOI:** 10.1101/2025.04.26.650728

**Authors:** David Wu, Liang Feng

**Author notes:** **Correspondence and requests for materials** should addressed to L.F.

## Abstract

Deep learning approaches, exemplified by AlphaFold, have made enormous progress in addressing the grand challenge of protein structure prediction. However, these methods typically predict only a single conformation, while accurately predicting multiple conformations, which are essential for the functions and regulations of many protein classes, remains challenging. Here, we demonstrate that AlphaFold2 has internal knowledge of alternative protein conformations, which can be effectively extracted by learning modifications to its intermediate hidden states using a negative entropy loss. Our resulting method, Entropy Guided Fold (EGF), achieves a success rate of over 80% in accurately predicting two conformations of 37 tested membrane proteins within a 2.0 Å RMSD threshold, with similar performance for proteins both within and outside AlphaFold2’s training dataset. In addition, our method successfully reveals three or more conformational states in several cases. We have also developed a post-processing strategy to effectively select accurate predictions. Overall, our method provides an efficient and robust approach for predicting multiple conformations of target proteins, with broad potential applications in mechanistic studies and translational applications.

## Introduction

The structure of a protein determines its function, and accurately predicting protein structure from sequence has been a long-standing challenge in biology. Recent breakthroughs in deep learning approaches such as AlphaFold^1^, RoseTTAFold^2^, and ESMFold^3^, have achieved atomic-level accuracy in predicting structures for a wide range of proteins. These tools have revolutionized research across nearly all areas of biological and biomedical science.

However, a significant portion of proteins such as transporters, receptors, enzymes, and regulatory proteins carry out their functions by transitioning between multiple conformations in response to external conditions, enabling diverse biological and physiological roles. While AlphaFold and related tools excel at predicting single conformations, they typically struggle to predict distinct conformations for proteins with multiple states^4-7^.

Recent innovations have sought to address this limitation by biasing predictions towards specific conformations, therefore expanding AlphaFold’s capabilities. These approaches primarily involve multiple sequence alignment (MSA) modification and subsampling or contact-based strategies. Given that AlphaFold2 (ref.^1^) relies on coevolutionary information from MSAs^1,7,8^ to infer residue-residue contacts in the 3D structures, modifying the MSA may influence the resulting AlphaFold2 prediction towards one conformation or another. Methods such as random sampling of shallow MSAs^7^, clustering and inputting MSA subsets^8^, and masking parts of MSAs to disrupt contacts (e.g., SPEACH_AF^9^) have been used to explore alternative conformations. Alternatively, contact-based methods have also been applied to guide AlphaFold2 toward specific conformations by introducing residue-residue contact constraints, either based on experimental information, such as leucine cross-linking (AlphaLink^10^), or deep learning-based contact map predictions^11^. Similarly, methods such as DEERFold^12^ use experimental distance information to guide AlphaFold2 in predicting alternative conformations, but obtaining such data is a significant undertaking that is time-consuming and often experimentally challenging. Template-based approaches, which select templates from similar proteins^13^ or those with specific properties^14^, and iterative scoring with evolutionary methods like MultiSFold^15^ have also been employed to predict multiple-conformational states. While these efforts have shown success in various cases, major challenges remain including low success rates for the majority of targets, difficulty in identifying faithful alternative conformations among large number of predictions, and variability in accuracy.

Here, we demonstrate that AlphaFold2 has internal knowledge of alternative conformational states during its prediction process. We then designed a method, Entropy Guided Fold (EGF), to extract this knowledge by modifying AlphaFold2’s intermediate hidden states using auxiliary losses to reduce its bias towards one conformation. Using a curated evaluation set of 37 membrane proteins (24 primary validation and 13 additional test proteins) with experimentally determined structures in multiple conformations, we show that EGF can effectively and accurately predict multiple conformations, achieving over 80% success rate in predicting two major conformations within a 2.0 Å RMSD threshold when using an ensemble of 10 pairs of predicted models. Furthermore, we demonstrate that even when reducing the ensemble to two pairs of predicted structures, EGF maintains a 69% success rate on the set of 24 validation proteins.

### Distogram-guided Prediction

Based on the prior work summarized above, we hypothesized that AlphaFold2 inherently contains representations of alternative conformations but is biased towards one conformation, masking other(s) and resulting in consistent prediction of a single conformation. This hypothesis is supported by the partial success of MSA subsampling methods: since separate clusters may encode information on specific conformations, the full MSA must contain the necessary information to predict all these conformations. We thus reasoned that a method utilizing all MSA sequences should be more accurate than approaches that discard large fractions of sequences, as different parts of an MSA may contribute to predicting diverse conformations and improving structural prediction accuracy.

We subsequently explored methods to reduce AlphaFold2’s bias towards a single conformational state to allow for the prediction of alternative conformations. Our approach leveraged AlphaFold2’s auxiliary predictions, such as distograms (representation showing probability distribution of distances between all residue pairs), which we found not only provide additional prediction information, but also can serve as signals to guide the prediction process. We found that applying a loss to the distogram and backpropagating the resulting signals to modify intermediate hidden states at the beginning of the AlphaFold2 network could mitigate bias towards one specific conformation. By disrupting signals favoring one state, this approach could shift AlphaFold2’s prediction toward alternative conformational state(s).

To evaluate the effectiveness of distogram-guided prediction, we initially tested a loss function that used ground truth distograms from distinct conformations to supervise predicted distograms, aiming to steer AlphaFold2 towards a specific ground truth conformation. As a proof of principle, we focused on membrane proteins, which constitute approximately 25-30% of the proteome and 60-70% of drug targets^16^. These proteins mediate information relay, material transfer, and signal transduction across membranes, processes that typically require conformational changes. Understanding multiple conformations of membrane proteins is crucial for advancing drug discovery and elucidating their physiological functions. Special emphasis was placed on membrane transporters, a major class of membrane proteins with over 1,000 members in the human genome. These transporters exhibit vastly diverse sequences, folds, and functions, playing vital roles in material exchange, signaling transduction, and serving as major drug targets for various diseases. Transporters undergo conformational transitions during their transport cycle to give rise to alternating-access transport process for substrate translocation across membranes. Understanding their major conformational states is crucial for elucidating transport mechanisms and designing selective therapeutic modulators. Decades of dedicated experimental efforts in the field have led to structural determination of a number of transporters in multiple conformational states, providing an ideal test case. Using a curated validation set of 24 membrane proteins—primarily transporters but also including enzymes—spanning diverse sequences and structures, we show that distogram-guided folding successfully guides AlphaFold2 to predict alternative conformations in all cases, achieving a significant mean reduction in RMSD toward the ground truth alternative conformation, even when AlphaFold2 initially exhibits a strong bias toward a different conformation (Figure 1a-c).

**Figure 1:**
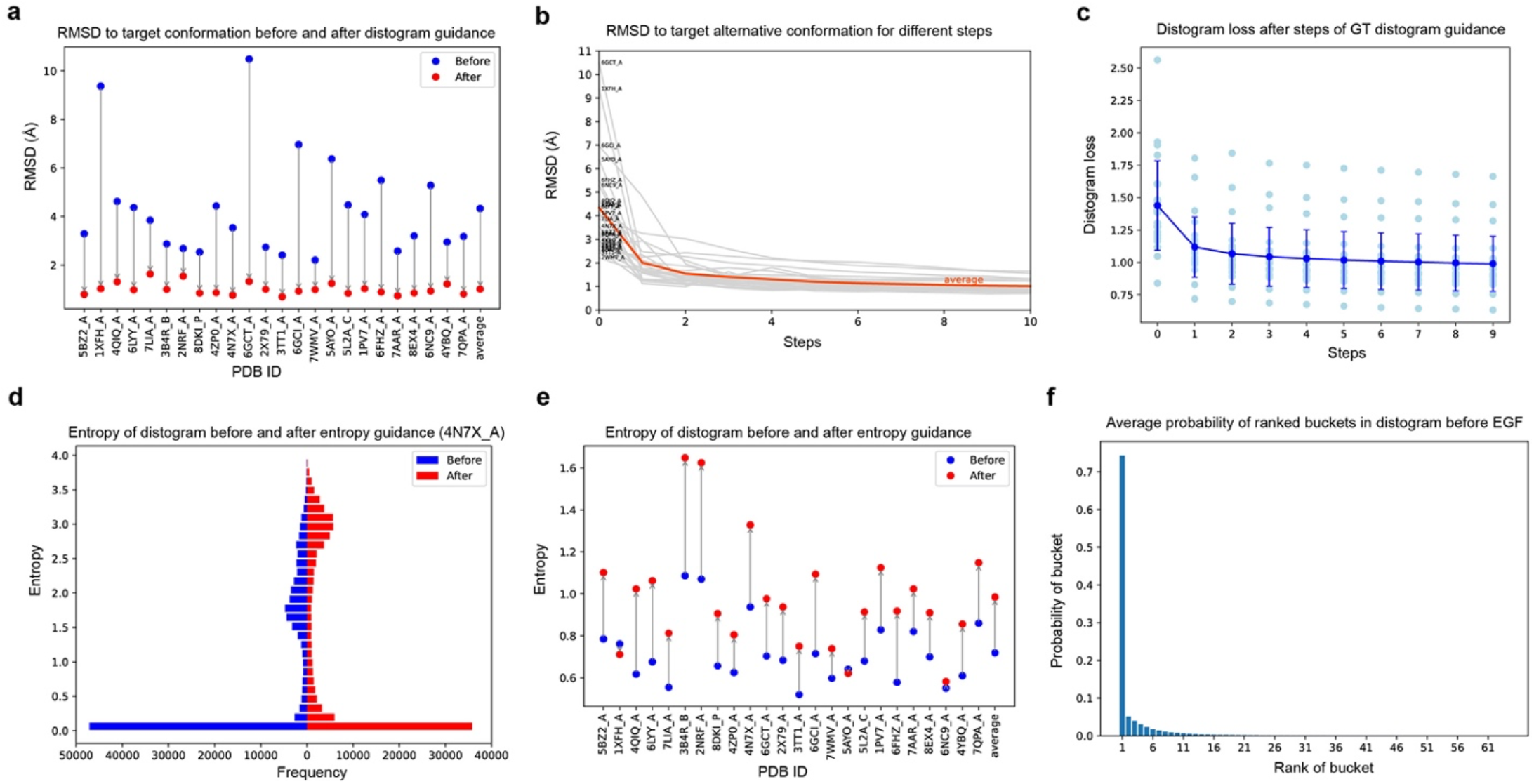
Analysis of distogram and entropy guidance. **Top row:** Results for distogram-guided folding using ground truth distograms on a set of 24 membrane proteins, using model 3 of AlphaFold2 for 10 distogram guidance steps. **a**, Reduction in RMSD (Å) to target state after applying distogram guidance for 10 steps. **b**, RMSD to target state after different numbers of distance guidance steps. **c**, Ground truth distogram loss decreases after applying ground truth (GT) distogram guidance, indicating that the probability mass is being transferred to the ground truth bucket. **Bottom row:** Entropy of 64-bin distograms before and after entropy guidance on a sample protein (PDB: 4N7X_A)^17^ and on all 24 membrane proteins. **d**, Back-to-back histogram comparing mean entropy of distograms before and after entropy guidance on 4N7X_A. **e**, Changes in mean entropy of distograms before and after entropy guidance for 24 membrane proteins. **f**, Average probability of ranked buckets over all 24 membrane proteins. Results show that the top bucket contains a large portion of the probability mass, indicating that the model is heavily biased towards one conformation.

### Entropy-Guided Prediction

Given these promising results, we shifted our focus to developing a loss function capable of effectively steering AlphaFold2 toward predicting alternative conformational states without relying on ground truth information, which is needed to enable *de novo* prediction of multiple conformational states.

To gain a better understanding of AlphaFold2’s bias and design an effective loss function, we examined the predicted distogram for a sample protein, 4N7X_A^17^. We found that the probability distributions at many points throughout the distogram were concentrated into a single bucket, indicating AlphaFold2’s strong bias towards one conformation (Figure 1f). In comparison, applying the ground truth distogram loss shifts the distogram to be concentrated at the ground truth alternative bucket as demonstrated by the decrease in distogram loss (Figure 1c). We hypothesized that introducing *uncertainty* in the distogram could effectively reduce AlphaFold2’s bias towards one conformation, thereby promoting exploration of alternative conformations. To achieve this, we designed a loss function (equal to the negative entropy of each residue pair’s distance distribution, averaged over all residue pairs) that incentivizes unbiased predictions by elevating uncertainty, measured by entropy^18^, over the distance buckets in the distogram, where *B* is the number of distance buckets, *N* is the number of residues in the protein, and *d* is the distogram indexed by the pair of residue indices and the bucket index:

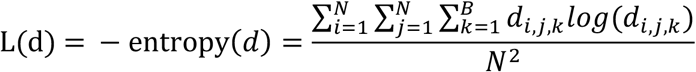

The resulting entropy augmentation is designed to reduce AlphaFold2’s bias towards a single conformation, thereby enabling prediction of alternative conformational states. The process begins by running the input sequences through AlphaFold2 to generate an initial predicted distogram and structure. The distogram signal is then backpropagated through the model to refine the input representations, incentivizing modifications that increase distogram entropy (for ease of implementation in deep learning frameworks, entropy augmentation is formulated as reducing the negative entropy). Finally, the refined representations are passed through AlphaFold2 to obtain a predicted structure that may correspond to an alternative conformation (Figure 2) (See Methods for details). We refer to this entropy-guided, distogram-driven method as *Entropy Guided Fold (EGF)*. Consistent with our hypothesis, entropy guidance increased the entropy of the distograms (Figure 1e). For example, using 4N7X_A as a representative case, entropy guidance shifted the probability distribution of the distogram entropy towards higher values (Figure 1d).

**Fig. 2:**
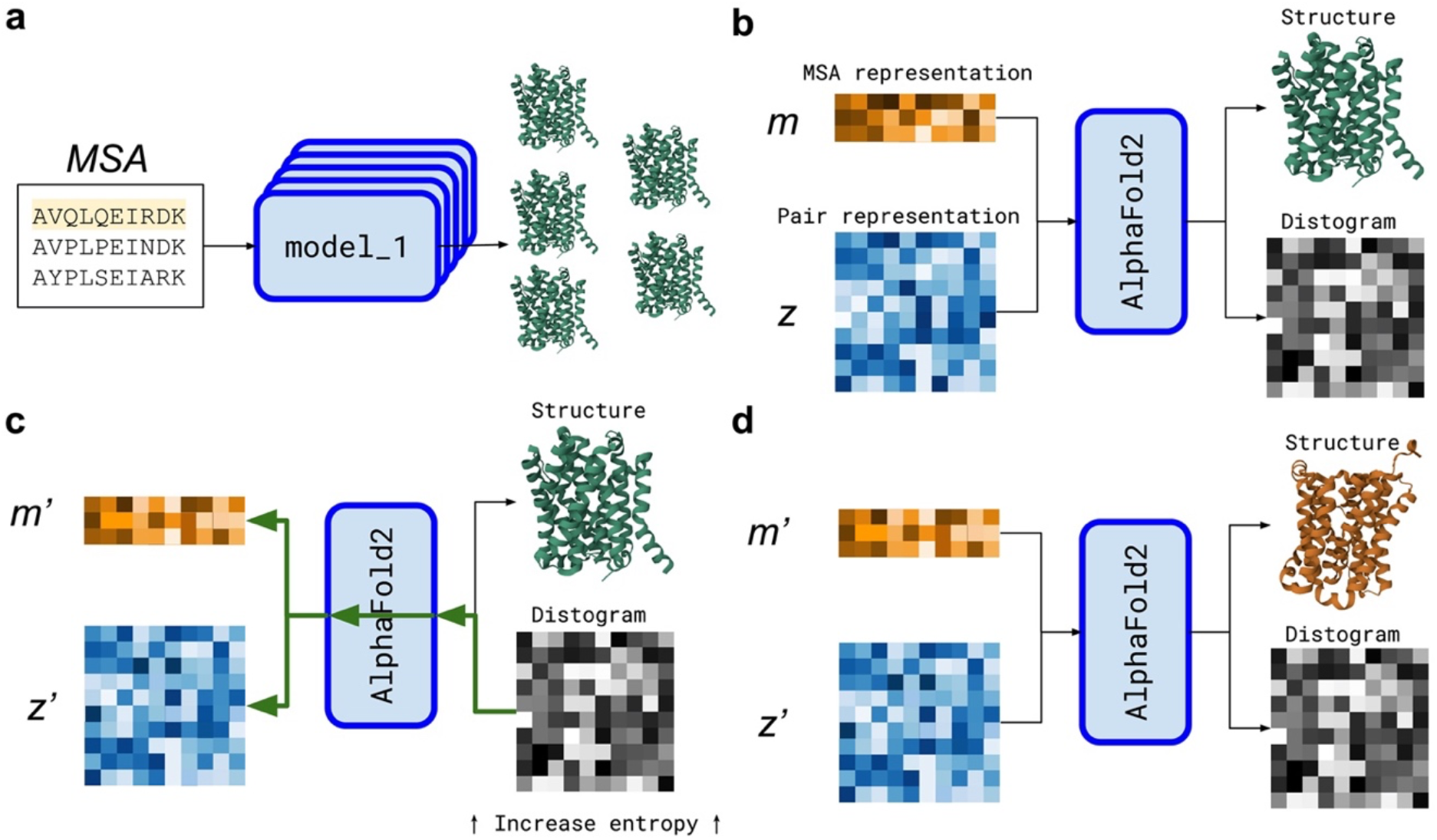
Illustration of Entropy Guided Structure Prediction. **a**, Running an ensemble of AlphaFold2 models on a given input protein typically leads to predictions of the same conformation. **b**, The first step of entropy guided structure prediction is the same as running the original AlphaFold2 model, using the input *m* and *z* representations to predict a structure and a distogram. **c**, Signal from the distogram is back propagated through the model to learn modifications to the *m* and *z* representations that lead to greater distogram entropy. **d**, The modified *m* and *z* representations are run through AlphaFold2 to obtain an alternative conformation.

To evaluate how increased distogram entropy impacts AlphaFold2’s ability to predict alternative conformations and assess whether it reduces bias towards a single conformation, we tested our EGF method on the same validation set of 24 cases. Notably, our method does not require training, so no training dataset was used. For each case, ten AlphaFold2 predictions were performed, generating two structures per prediction (with or without one step of EGF), and a total of 20 predicted structures per protein target. We evaluated variants of the entropy guided prediction by varying learning rates (0.001, 0.01, or 0.1) and step numbers and selected the best-performing variant, which had a learning rate of 0.01 and one step of EGF. Experimental conformations for each target were identified through a PDB database search and compared with the predicted structures from EGF. Our results show that, with EGF, AlphaFold2 successfully predicted two main known conformations in the majority of cases. Notably, 20 out of 24 cases produced predictions of both conformations that closely matched the experimental structures, with a Cα RMSD below a 2.0 Å cutoff (Table 1). Importantly, the prediction accuracy was consistent regardless of whether the experimental structures were deposited before or after the AlphaFold2 training date cutoff (Table 1), indicating that EGF-enabled predictions do not rely on prior knowledge from AlphaFold’s training data.

**Table 1:** Comparison of predicted structures from EGF with experimental structures. For each target protein, the PDB IDs of two distinct conformations are provided, along with the closest predicted structure for each conformation. For the best prediction corresponding to each conformation, the RMSD to both ground truth conformations is reported in angstroms, formatted as (RMSD to conformation #1, RMSD to conformation #2). Notes on specific sequences used are provided in Methods.

| Before AlphaFold2 training date cutoff |  |  |  | After AlphaFold2 training date cutoff |  |  |  |
| --- | --- | --- | --- | --- | --- | --- | --- |
| PDB #1 | Best #1 (Å) | Best #2 (Å) | PDB #2 | PDB #1 | Best #1 (Å) | Best #2 (Å) | PDB #2 |
| 4BWZ_A | **0.87**, 3.40 | 4.22, **1.36** | 5BZ2_A | 7CKO_A | **1.03**, 3.70 | 3.57, **1.30** | 7CKR_A |
| 6X16_C | **0.97**, 9.39 | 9.48, **1.12** | 6BMI_B | 7TXT_S | **0.87**, 3.75 | 1.84, 3.22 | 6DZZ_A |
| 4JA4_C | **1.13**, 4.57 | 3.76, 2.36 | 6N3I_A | 8DYP_A | **1.06**, 3.33 | 3.40, **1.35** | 8DKX_P |
| 3B4R_A | **1.50**, 3.09 | 3.28, **1.77** | 3B4R_B | 6EUQ_A | **0.56**, 4.43 | 2.62, 2.71 | 6GV1_A |
| 6PJU_A | **0.59**, 2.69 | 1.09, 2.18 | 2NRF_A | 8OUJ_B | **1.71**, 10.79 | 9.15, **1.91** | 8OUJ_C |
| 7CYK_B | **0.63**, 3.38 | 3.37, **0.90** | 6LH1_A | 4C9G_A | **1.05**, 6.71 | 5.87, **1.88** | 6GCI_A |
| 4D1C_A | **1.27**, 2.75 | 2.89, **1.74** | 2X79_A | 7WMV_A | **1.22**, 2.24 | 2.86, **1.63** | 7SLA_A |
| 7DIX_A | **0.66**, 2.43 | 2.04, **1.00** | 6XWM_A | 3VVN_A | **0.62**, 5.27 | 3.80, **1.96** | 6FHZ_A |
| 5AYM_A | **1.11**, 6.04 | 6.14, **1.52** | 6BTX_A | 7AAR_A | **1.79**, 3.25 | 3.03, **1.99** | 6H7D_A |
| 5L2B_A | **0.92**, 4.94 | 4.35, **1.27** | 5L2B_C | 8EX6_A | **0.87**, 3.90 | 4.01, **1.01** | 8EX7_A |
| 5GXB_A | **1.63**, 4.18 | 4.34, **1.69** | 2Y5Y_A | 5T77_A | **1.49**, 5.78 | 6.13, **1.74** | 6NC9_A |
| 4YB9_D | **1.49**, 4.26 | 3.59, **1.75** | 4YBQ_A | 7QPC_B | **1.29**, 3.02 | 3.18, **1.40** | 7QP9_B |

In contrast, without EGF, AlphaFold2 predominantly predicted only one conformation. In fewer than 5 out of 24 cases (Table 2), AlphaFold2 produced predictions matching both conformations across 20 predictions. The recently developed AF-Cluster method^8^ markedly improved prediction efficiency, successfully predicting both conformations in 9 out of 24 (38%) cases. In comparison, our EGF-based method substantially improves upon the current state-of-the-art, achieving successful predictions of both conformations with high accuracy in 83% of tested cases, demonstrating its feasibility for reliably obtaining multiple conformations.

**Table 2:** Proportion of target proteins for which both conformations are successfully predicted within an RMSD threshold of 2.0 Å. Target proteins with both experimental structures released before the AlphaFold2 training date cutoff are listed separately from those with at least one experimental structure released after the cutoff. Each cell shows mean proportion ± the sample standard deviation across three seeds (only one seed is run for AF-Cluster due to computational cost).

| Method | Before AF2 train cutoff | After AF2 train cutoff | Total |
| --- | --- | --- | --- |
| AF-Cluster | 5/12 = 0.417 | 4/12 = 0.333 | 9/24 = 0.375 |
| AF2 x 10 models | 0.167 ± 0.000 | 0.222 ± 0.048 | 0.194 ± 0.024 |
| EGF x 10 models | **0.861 ± 0.048** | **0.806 ± 0.048** | **0.833 ± 0.042** |

Consistent with the low RMSD values, the successfully predicted structures superimpose very well onto the experimental structures of both conformations, respectively, as illustrated by examples in Figure 3. Each successfully predicted conformation accurately reflects the corresponding experimental conformation and is clearly distinct from the alternative conformation. In many cases, multiple closely related predictions correspond to a specific experimentally-determined conformation. Notably, the successful predictions match nearly perfectly with the ordered secondary structural elements of the experimental structures. Deviations are primarily observed in loops connecting structured elements, which are often flexible. These observations suggest that the EGF-facilitated predictions faithfully model the alternative conformations of the target proteins.

**Figure 3:**
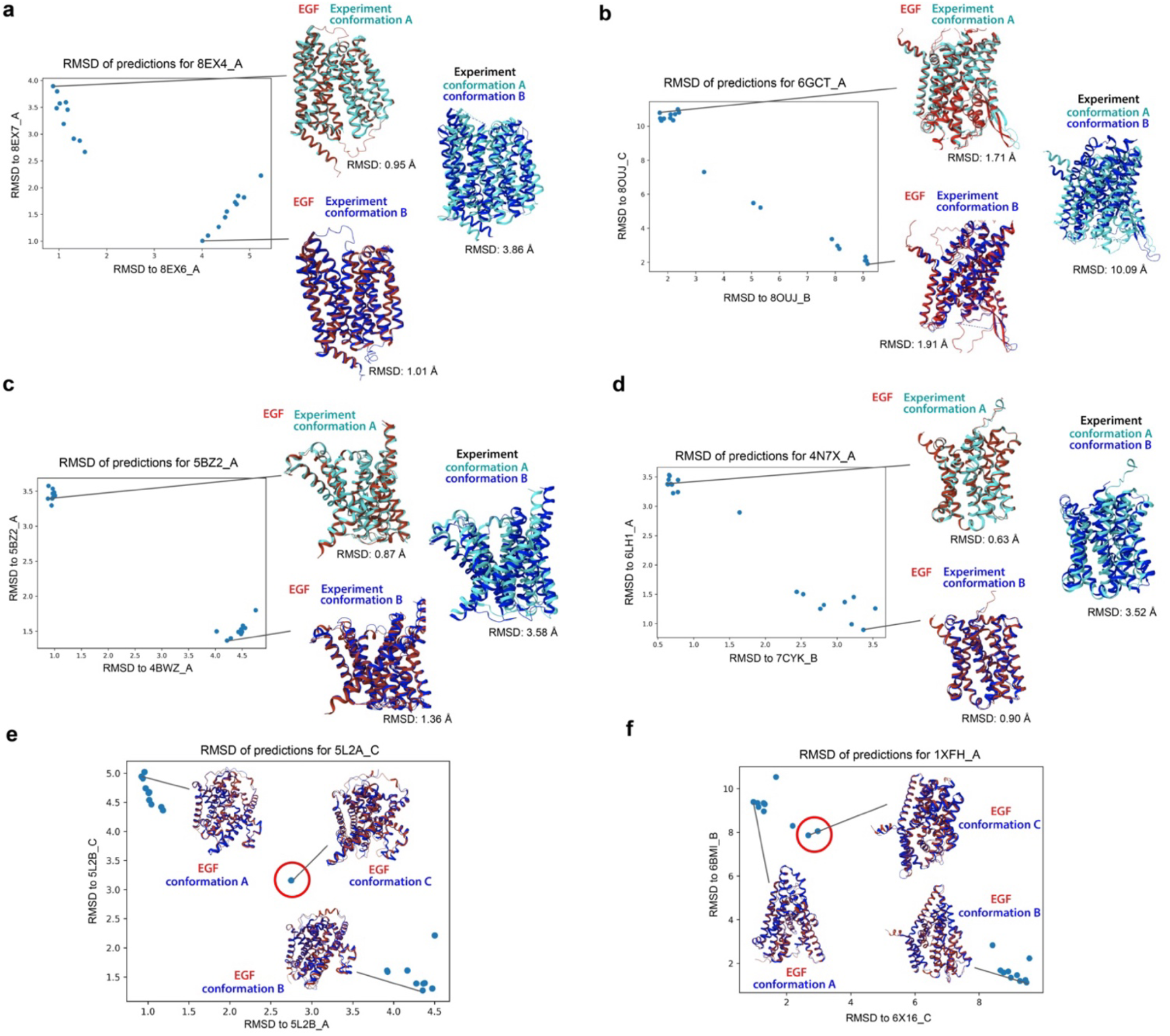
Comparison of the EGF predicted structures with the experimental structures. **a-d**, Comparison of predicted and experimental structures for both conformations. RMSD plots of predicted structures against both conformations are shown for 8EX4_A^19^, 3B4R_B^20^, 5BZ2_A^21^, and 4N7X_A^17^ (from seed 0). Predicted structures are colored in red, while experimental structures are colored in cyan and blue. Superimposition of experimental structures in both conformations are also shown. **e-f**, EGF predictions match more than two conformational states of experimental structures. Red-circled points highlight predictions corresponding to intermediate states, in addition to the two primary conformations represented on the axes.

### Relationship between negative entropy loss and ground truth distogram loss

To assess the relationship between negative entropy loss and ground truth distogram loss in facilitating alternative conformation predictions, we measured the ground truth distogram loss while only optimizing negative entropy loss. Our analyses revealed that while ground truth distogram loss to the alternative conformational state tends to decrease during the first optimization step of entropy loss (Extended Data Fig. 1), additional steps lead to an increase in ground truth distogram loss despite a continued decrease in negative entropy loss, causing predicted structures to diverge away from the ground truth structures. These findings suggest that applying entropy guidance for one step strikes the most effective balance between increasing uncertainty and preserving AlphaFold2’s internal knowledge of an alternative conformation.

### Predicting more than two conformations

The results summarized in Tables 1 and 2 are based on comparing predictions with two experimentally determined ground truth conformations. However, biophysical and structural studies have demonstrated that many proteins sample multiple discrete conformational states. For example, membrane transporters typically cycle between inward-open, occluded, and outward-open conformations during a transport cycle. We hypothesized that some predictions from entropy-guided prediction, which do not correspond to the two given conformational states, might represent additional conformational states of the target proteins.

Indeed, we found that several predictions matched additional experimentally determined conformational states. For example, in a nucleoside transporter, while most predictions from entropy-guided prediction correspond to either the inward-open or outward-open states, one prediction, which deviates significantly from both states, matches well to an intermediate state (PDB: 5L24_C^22^; RMSD= 1.79 Å) (Fig. 3e). Similarly, for a glutamate transporter, while most predictions corresponded to either the inward- or outward-facing states (PDB: 6X16_A^23^ and 1XFH_A^24^, respectively), we found predictions that match the intermediate chloride-conducting state (PBD: 6WYK_A^25^; RMSD= 1.78 Å) (Fig. 3f). These findings demonstrate that entropy-guided prediction is capable of predicting biologically relevant, intermediate conformational states.

Due to the challenges of capturing experimental structures for multiple conformational states of a target protein, the majority of proteins in the Protein Data Bank still lack structures beyond a single conformation, and it is even rarer to find cases with more than two states, despite their existence often being suggested by other studies. We suspect that, for several target proteins, predictions that do not correspond to a known experimentally described conformational state may potentially represent biologically relevant conformations that have yet to be experimentally determined. For example, in the case of a multidrug efflux transporter, one prediction that does not match either the ground truth inward-open or the outward-open states (PDBs: 6EUQ_A^26^ and 6GV1_A^27^) instead exhibits characteristics of an occluded state (Figure 4a), where both the outer and inner gates close around the central cavity. Similarly, for a ferroportin family iron transporter, two predictions form a separate cluster distinct from the two experimentally determined conformational states (PDBs: 5AYM_A^28^ and 6BTX_A^29^) (Figure 4b). Upon manual inspection, these predictions appear to represent a partially inward-facing state, where the inner gate partially closes. These findings suggest that entropy-guided prediction may potentially predict novel conformations. Future experimental work will be needed to validate these predictions for target proteins of interest.

**Figure 4:**
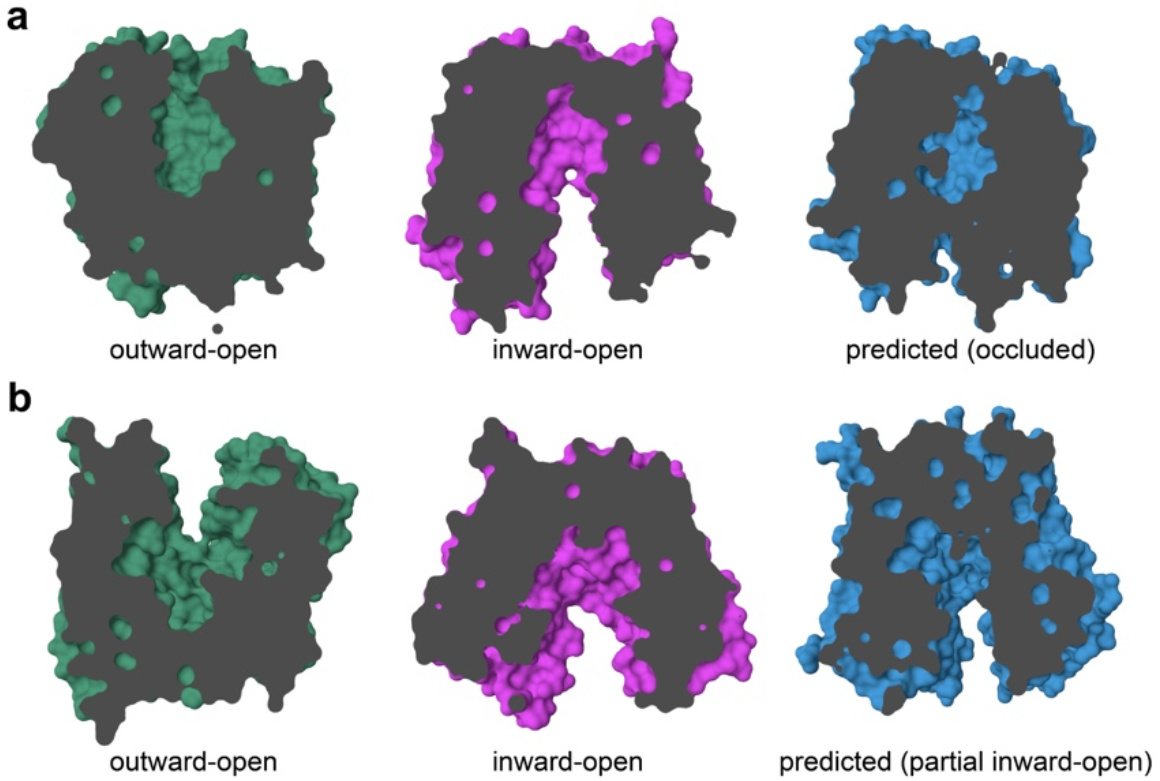
The potential prediction of novel conformations. **a**, The predicted structure of a multidrug efflux transporter MdfA (right) is compared with experimental structures (outward-open, PDB: 6EUQ_A^26^, left; inward-open, PDB: 6GV1_A^27^, center). The predicted structure on the right appears to represent an occluded conformation. **b**, The predicted structure of an iron transporter ferroportin is compared with experimental structures. Experimental structures of the outward-open (PDB: 5AYM_A^28^) and inward-open (6BTX_A^29^) conformations are shown on the left and center, respectively. The predicted structure on the right appears to represent a partially inward-facing conformation.

### Prediction clustering and intra-cluster variation

Among the 20 EGF-predicted structures of a given query protein, there are typically multiple predicted structures resembling a specific known conformation. In the majority of cases, the predicted structures can be grouped into a small number of defined clusters (Figure 5a & b, Extended Data Table 1) as assessed by DBSCAN clustering^30^, which estimates groups of similar predictions based on density. To reduce the likelihood of random outliers, we focused only on groups containing more than two members. Overall, successful predictions of distinct conformations generally fall into discrete clusters. The observed small number of clusters is in line with the idea that many proteins may adopt a set of relatively stable yet distinct conformational states. In some cases, no clearly defined clusters were observed; instead, the predictions formed a linear distribution pattern between the two ground truth conformations and were either grouped into one group or split into many small groups, which did not have enough members to be counted as clusters (Figure 5c). This raises an intriguing question of whether, for these proteins, multiple intermediate conformations may bridge the two primary conformations.

**Figure 5:**
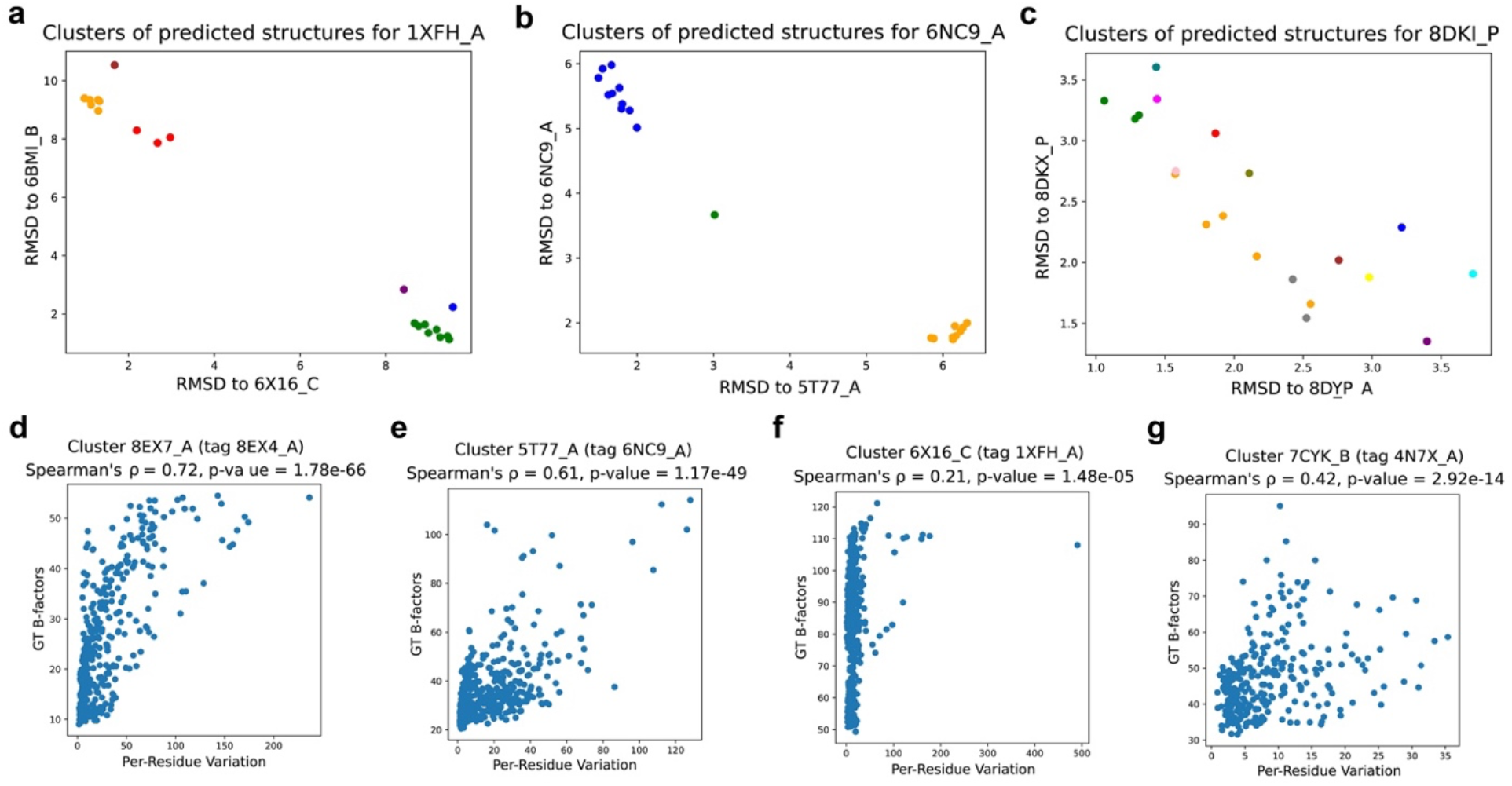
Clustering of predicted structures. **a-b**, Example clusters of predicted structures for 1XFH_A^24^ and 6NC9_A^31^. Each color represents a distinct group by DBSCAN. **c**, Example of a protein where predictions form a linear pattern between the two ground truth conformations. **d-g**, The correlation between the experimental B-factors and the computed per-residue variation of the predicted structures within a cluster. Only clusters with more than two members are analyzed. Four clusters, each clearly corresponding to a ground truth conformation are shown here as samples. A positive correlation was observed in these cases, as indicated by the low p-values from Spearman’s test.

Within a cluster, the predicted structures typically correspond to the same ground truth conformation. Interestingly, we found that per-residue variation among predicted structures within a cluster positively correlates with the experimental B-factors from the PDB file of the closest ground truth conformation (Figure 5d-g). Specifically, we analyzed clusters that are within 2.0 Å of one (but not both) ground truth conformations. Among 37 resulting clusters, the mean Spearman correlation was 0.316. Notably, 31 of the 37 clusters had a

Spearman correlation of greater than 0.1, all with p-values below 0.05. This correlation between the variation in residue coordinates of predicted structures and the experimental B factors, which generally reflect flexibility/mobility of a residue, raises the possibility that the variations in the predicted structures may potentially reflect, to some degree, the local conformational dynamics of the query protein. The biological relevance of these structural variations in the predicted structures warrants future investigation through biophysical and structural studies.

### Post-processing

Although generating 20 predicted structures per target protein increases the probability that both ground truth conformations emerge from the predictions, the uncertainties in *which* predicted structures most likely correspond to a ground truth conformation complicate the practical utility of the predictions.

To alleviate this problem, we examined various ranking and scoring methods. First, we used AlphaFold2’s built-in pLDDT ranking function with a mean taken over all residues as a ranking score. We found that pLDDT was heavily skewed toward the original AlphaFold2 predicted structures (before applying EGF) and showed significantly lower scores for structures that matched alternative conformations obtained after applying EGF (Figure 6a). Furthermore, we found that pLDDT scores did not indicate closeness to a conformational state even within the set of structures from before or after applying EGF. Therefore, pLDDT was not a good indicator of closeness to ground truth conformations.

**Figure 6:**
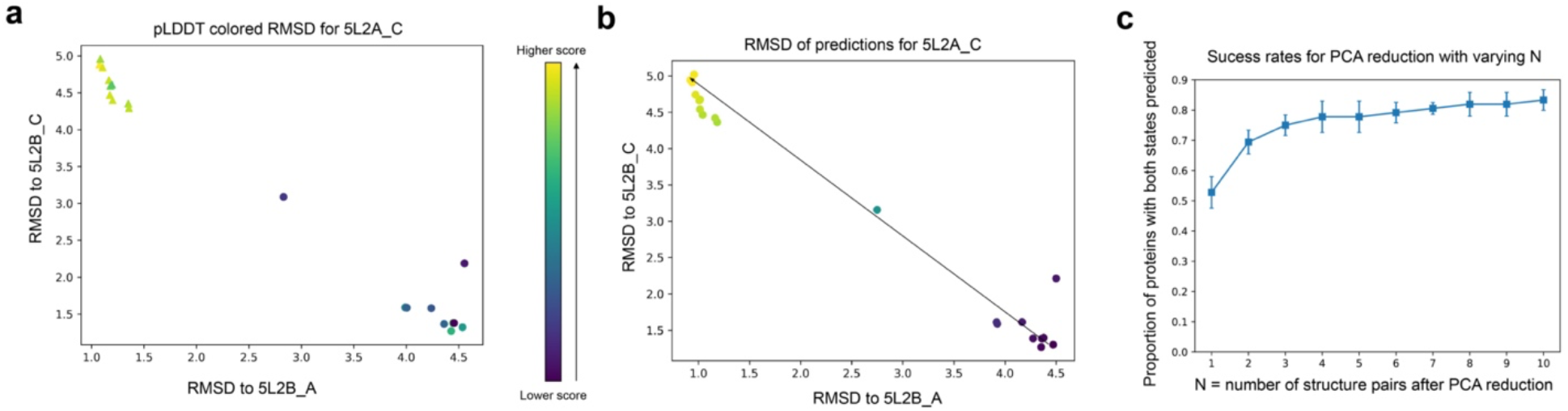
pLDDT vs PCA scoring. **a**, pLDDT score does not directly measure similarity to ground truth conformational states. Predicted structures generated before applying EGF (triangles) have significantly higher pLDDT scores than those generated after applying EGF (circles). All points are colored using the corresponding score. **b**, Using PCA dimension reduction, the predicted structures form a linear trajectory connecting one ground truth conformation to the other. **c**, The proportion of proteins with both conformations successfully predicted from the test dataset with 24 targets with multi-conformations was plotted against the number of output structures after PCA reduction. Each point represents the mean proportion ± sample standard deviation across seeds 0, 1, and 2.

Instead, we found that for the majority of proteins, applying PCA dimensionality reduction to the raw 3D coordinates and reducing each structure to a single numerical value revealed a strong linear pattern along the direction between the two ground truth conformations (Figure 6b). In a small fraction of cases, the pattern was more complex (Extended Data Fig. 2). Notably, by selecting structures with the highest and lowest PCA scores, much of the performance of the full 20-structure ensemble can be recovered using significantly fewer structures (Figure 6c and Extended Data Table 2). For example, selecting the top three pairs of predictions successfully identified two major conformations (RMSD < 2.0 Å) with a success rate of 0.750 (~ 18 out of 24 cases). Even with just the top predicted pair, this approach successfully predicted both conformations in more than half of the cases (53%).

### Additional test set of membrane proteins

To further evaluate the performance of the optimized EGF method, we curated an additional set of membrane proteins whose structures were only recently released, all postdating the AlphaFold2 training cutoff^1^. These proteins were selected after the establishment of the EGF method and were not used for evaluating variants with different hyperparameters. Using this test set, we found the EGF-facilitated method successfully predicted two major conformations in 10 out of 13 cases (Tables 3 & 4), applying the same RMSD cutoff of 2.0 Å. This success rate is comparable to our evaluation set, where the expected success rate would have been 11 out of 13, indicating statistically similar performance.

**Table 3:** Comparison of predicted structures from entropy-guided prediction with experimental structures for a new test set of target proteins. For each target protein, the PDB IDs of two distinct conformations are provided, along with the closest predicted structure for each conformation. For the best prediction corresponding to each conformation, the RMSD to both ground truth conformations is reported in angstroms, formatted as (RMSD to conformation #1, RMSD to conformation #2). Notes on specific sequences used are provided in methods section.

| Additional test set of target proteins |  |  |  |  |  |  |  |  |
| --- | --- | --- | --- | --- | --- | --- | --- | --- |
| PDB 1 | PDB 2 | Best 1 (Å) | Best 2 (Å) |  | PDB 1 | PDB 2 | Best 1 (Å) | Best 2 (Å) |
| 8VZN_A | 8VZO_A | **0.67**, 3.48 | 3.02, **1.57** |  | 7PQG_A | 7VAG_A | **1.84**, 4.04 | 5.07, 2.27 |
| 8WNY_B | 8WNT_B | **0.96**, 2.19 | 1.59, **1.45** |  | 8JZU_A | 8P6A_A | **1.52**, 4.38 | 4.92, **1.88** |
| 8J74_A | 8J76_A | **1.38**, 2.67 | 2.95, **1.61** |  | 8SC3_A | 8JTZ_A | **1.10**, 3.27 | 2.82, **1.77** |
| 8WFK_A | 6ZPL_A | **1.26**, 2.67 | 2.15, **1.68** |  | 8WBX_B | 8I3B_B | **1.58**, 3.98 | 2.75, 2.52 |
| 8WM7_A | 8W9M_A | **1.10**, 2.60 | 1.15, 2.60 |  | 8J7X_B | 8J7W_B | **1.69**, 3.17 | 4.14, **1.87** |
| 8JT5_A | 8X09_A | **1.22**, 3.31 | 3.52, **1.45** |  | 8W6T_A | 8W6T_B | **1.79**, 5.46 | 6.96, **1.07** |
| 8TGI_A | 8TGG_B | **0.91**, 3.52 | 3.24, **1.49** |  |  |  |  |  |

**Table 4:** Proportion of target proteins for which both conformations are successfully predicted within an RMSD threshold 2.0 Å, on a test set of 13 membrane proteins. Each cell lists mean proportion ± sample standard deviation of proportion across three seeds.

| Method | Additional test set of 13 target proteins |
| --- | --- |
| AF2 x 10 models | 0.128 ± 0.044 |
| EGF x 10 models | **0.743 ± 0.044** |

## Discussion

Deep learning-based methods have delivered remarkable success in predicting protein structures in single conformations, but predicting multiple conformations remains a considerable challenge. Experimentally capturing alternative conformations is typically challenging and often relies on fortuitous discoveries, as no standardized methods exist to stabilize proteins in specific conformations. To address this gap, we hypothesized that AlphaFold inherently has the capacity to predict multiple conformations, despite its default algorithm strongly favoring a single state. To assess this, we developed a distogram-guided method that incorporates a negative entropy loss to incentivize uncertainties in the distogram, which reduces AlphaFold’s bias toward a single conformation to allow the exploration of alternate conformations. This approach achieved over 80% success rate in accurately predicting both conformations across 37 tested membrane proteins (combination of both sets) compared to known experimental structures. The actual success rate is likely even higher, as experimental structures for some conformational states are missing, and certain unmatched predictions clearly show characteristics of alternative conformations. By projecting the predicted structures onto a linear scale and selecting the structures at the extremes of the scale, we recovered much of the ensemble performance while reducing the predicted structure set to one or two pairs. Unlike input subsampling or modification methods, our approach allows AlphaFold2 to leverage the full MSA, enhancing accuracy of predicted structures. It is also significantly more efficient, requiring only 1-10 runs of AlphaFold2 rather than hundreds for subsampled MSAs and can finish within minutes on a consumer GPU.

Our results showed that the negative entropy loss effectively guides AlphaFold2 to predict alternative conformations, supporting the hypothesis that AlphaFold2, with its complete set of inputs, inherently contains information about alternative conformational states. By introducing uncertainty into the predicted distogram, entropy-guided prediction provides an effective method for extracting such information. In essence, information about conformation-specific protein contacts is embedded in the MSA, as the distinct conformational states critical to protein function likely present across homologues. These conformation-specific contact constraints are inherently incompatible with alternative conformations, serving as key determinants for distinct conformational states. Importantly, this ability to predict alternative conformations does not come from memorizing training structures but from AlphaFold’s capability to identify alternative conformations encoded in the MSA. This conclusion is further supported by the method’s equally robust performance on proteins outside AlphaFold2’s training dataset.

Notably, our EGF method also correctly predicts three or potentially more conformations in several cases, and positive correlations were observed between structural variations within predicted clusters and the B-factors of experimental structures. These raise the possibility that predictions deviating from the ground truth experimental structures may, to some extent, reflect conformational dynamics, intermediate conformations, or transient in-pathway states. Future computational and experimental studies are needed to probe these possibilities. If validated, EGF-facilitated predictions could further enrich our understanding of protein conformational plasticity and dynamics. Furthermore, in principle, EGF could be extended to other structure prediction methods and applied to the analysis of protein complexes. In addition, our method provides a foundation for future improvements, such as testing alternative divergences to entropy and progressively refining signal extraction, to more effectively and confidently predict multiple conformational states.

Overall, our EGF method provides a robust, efficient, and accurate approach for predicting protein structures in multiple conformational states. The ability to do so will foreseeably benefit studies of protein function across physiological and disease states, inform protein engineering strategies to stabilize specific conformations, and reveal conformation-specific structural features that can be targeted for therapeutic and biotechnological applications.

---

**Extended Data Figure 1:**
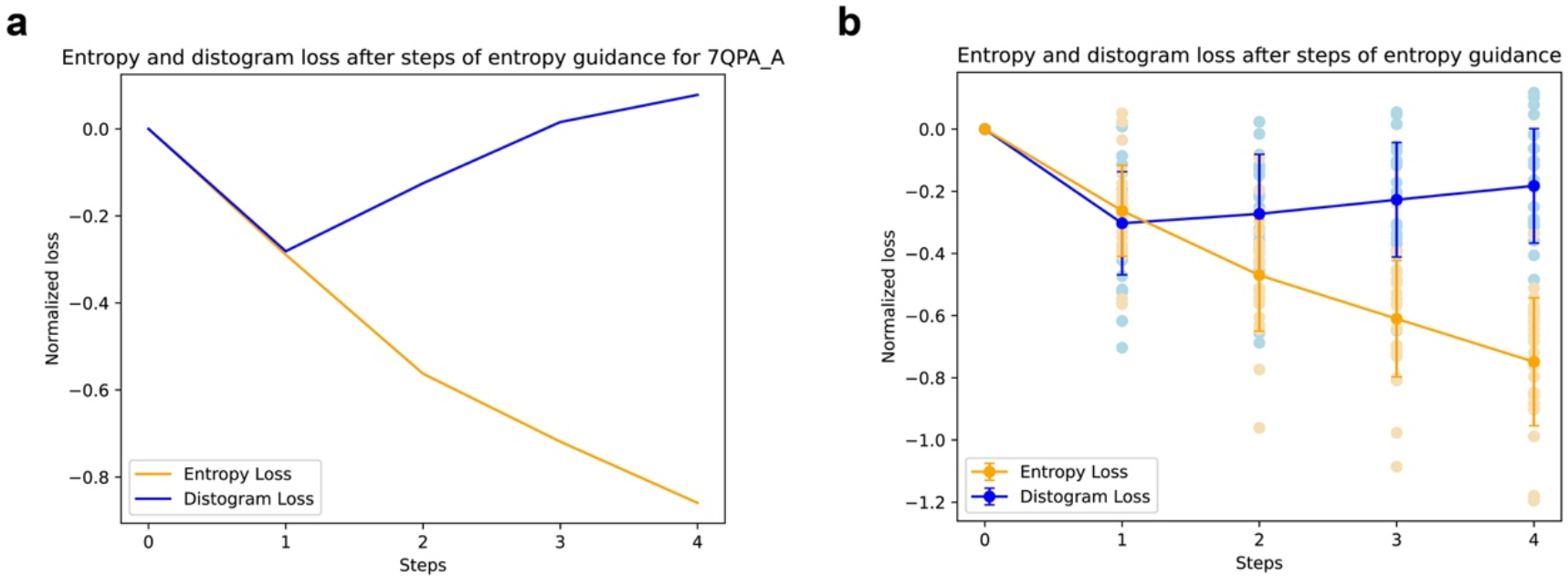
Relationship between distogram loss and negative entropy loss. Evaluation of the entropy and distogram loss after the indicated numbers of steps of entropy-guided prediction. To better highlight loss changes, we normalized losses by subtracting the first step loss. **a**, The negative entropy loss and the distogram loss for 7QPA_A over 5 steps. **b**, The negative entropy loss and the distogram loss for all 24 test proteins over five steps. Points represent individual values; the lines with error bars are mean and standard deviation, respectively.

**Extended Data Fig. 2:**
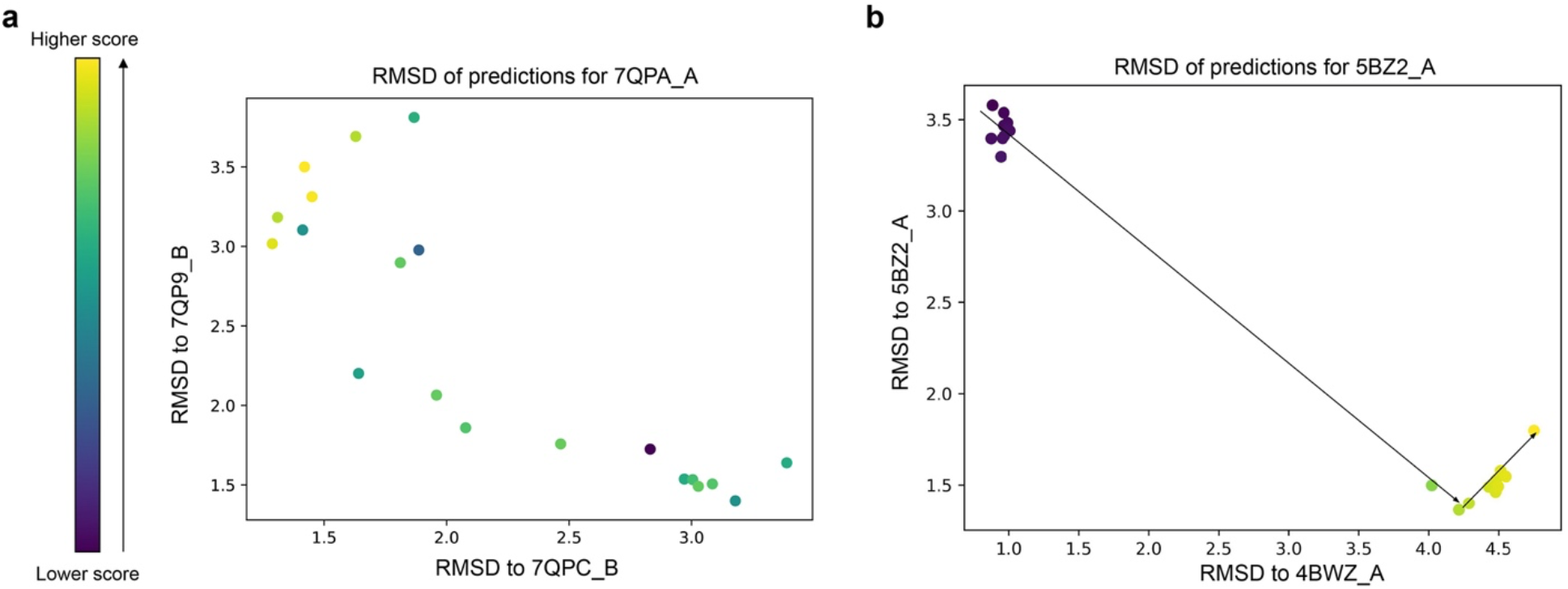
Examples of PCA scoring with more complex patterns in a small fraction of cases. **a**, For some proteins (e.g. 7QPA_A/7QPC_A), the PCA scores do not form a clear linear trajectory between two conformations, though a weak directional trend is observed. **b**, For certain proteins (e.g. 5BZ2_A/5BZ3_A), the PCA score-labeled predicted structures exhibit a hook-like pattern. In this pattern, after moving from one predicted structure to the other, some predicted structures go *past* the other ground truth conformation and begin moving away from it, which may potentially indicate an additional conformational state.

**Extended Data Table 1:** Cluster counts. A table summarizing the number of proteins with 1, 2, and 3 clusters. Most proteins have 2 clusters, while some have 1 cluster (indicating either unsuccessful prediction of multiple conformations or inability to resolve distinct clusters), and a few have 3 clusters (indicating the prediction of intermediate conformations). Only clusters with more than two members are counted.

|  |  |  |  |
| --- | --- | --- | --- |
| Clusters | 1 | 2 | 3 |
| # of proteins | 6 | 14 | 4 |

**Extended Data Table 2:** Proportion of proteins with both conformations successfully predicted under various PCA-based reductions using an RMSD threshold of 2.0 Å. Target proteins are categorized based on whether their structures become available before or after the AlphaFold2 training cutoff date. Each cell lists the mean proportion ± standard deviation across seeds 0, 1, and 2 (only one seed is run for AF-Cluster).

| Method | Before AF2 train cutoff | After AF2 train cutoff | Total |
| --- | --- | --- | --- |
| AF-Cluster | 5/12 = 0.417 | 4/12 = 0.333 | 9/24 = 0.375 |
| AF2 x 10 models | 0.167 ± 0.000 | 0.222 ± 0.048 | 0.194 ± 0.024 |
| EGF x 10 models | 0.861 ± 0.048 | 0.806 ± 0.048 | 0.833 ± 0.042 |
| PCA reduce to 3+3 | 0.805 ± 0.048 | 0.694 ± 0.048 | 0.750 ± 0.042 |
| PCA reduce to 2+2 | 0.722 ± 0.096 | 0.667 ± 0.000 | 0.694 ± 0.048 |
| PCA reduce to 1+1 | 0.583 ± 0.084 | 0.472 ± 0.048 | 0.528 ± 0.064 |

## METHODS

### Overview of AlphaFold2’s prediction algorithm

AlphaFold2 relies on three main sources of input: (1) the input protein sequence, (2) the input multiple sequence alignment (MSA), and (3) input structure template. These three inputs are transformed to obtain intermediate hidden states and passed through an evoformer module and a structure module to obtain a final predicted structure.

In addition to predicting the final protein structure, AlphaFold2 also makes several auxiliary predictions, including pLDDT, a predicted confidence metric that has been shown to provide an accurate estimate of protein structure accuracy, and a predicted distogram, which predicts the probability distribution of distances between all residue pairs in the final structure^1^.

Below, we provide an overview of AlphaFold2’s prediction algorithm relevant to our work. We remove templates and recycling as they are not used in this work.

#### Algorithm 1

AlphaFold2 inference without templates or recycling

Inputs: sequence *s*, MSA *a*

Outputs: Predicted structure *x*, Distogram *d*

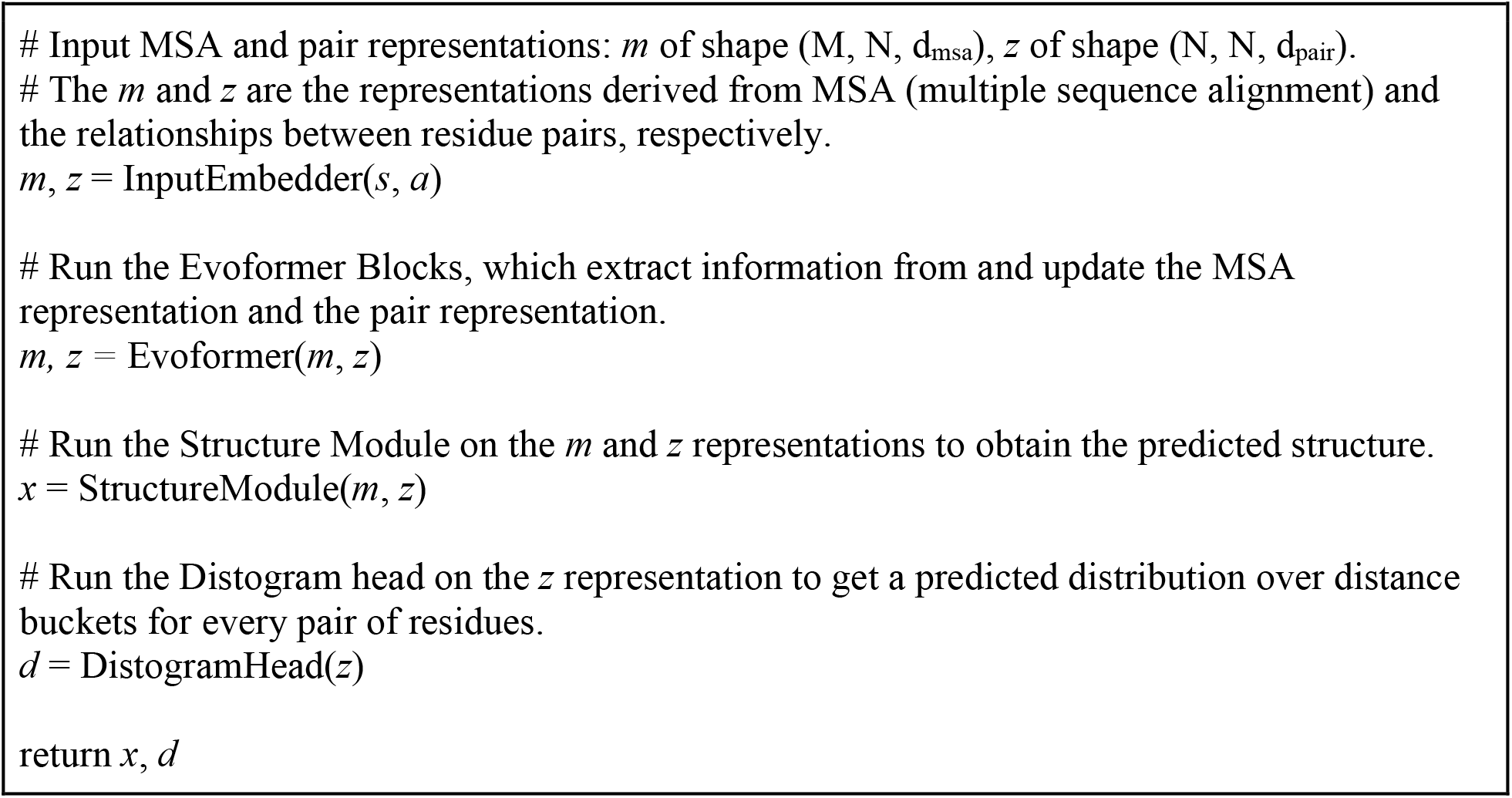

### Distogram-guided prediction algorithm overview

#### Algorithm 2

Entropy Guided Structure Prediction without templates or recycling.

Inputs: sequence *s*, MSA *a*, number of gradient steps G, LossFunction

Outputs: Predicted stru

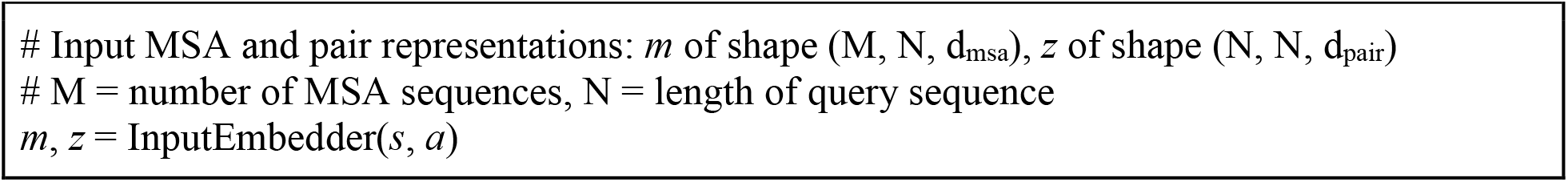
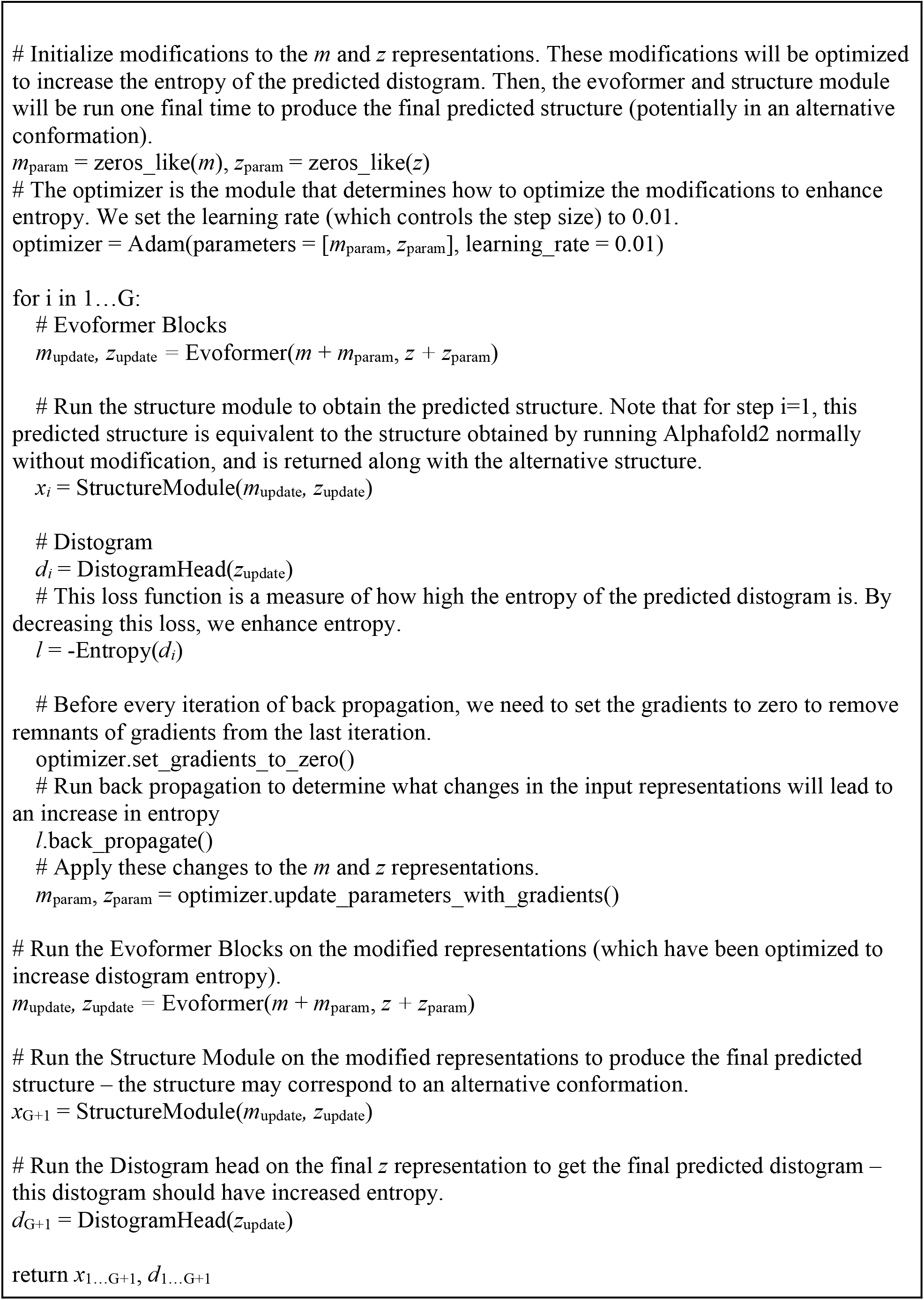

### Overview of PCA Reduction Method

After using AlphaFold2 and Entropy Guided Fold to obtain a set of 20 predicted structures (2 predicted structures for each of 10 models), we can use PCA reduction to reduce the number of predicted structures that most effectively capture distinct conformational states.

Specifically, our PCA reduction method takes the input structures, aligns them all to one structure using the Kabsch algorithm^32-34^, and uses sklearn PCA reduction^35^ (https://scikit-learn.org/stable/modules/generated/sklearn.decomposition.PCA.html) to reduce the raw 3D coordinates to a single number. The resulting PCA scores are sorted and the structures corresponding to the most extreme scores are selected and returned.

#### Algorithm 3

PCA Reduction of Entropy Guided Fold Predicted Structures. Inputs: Predicted structures *x*_1…N_, desired number of output pairs P Outputs: Reduced set of predicted structures *x*_1…P_

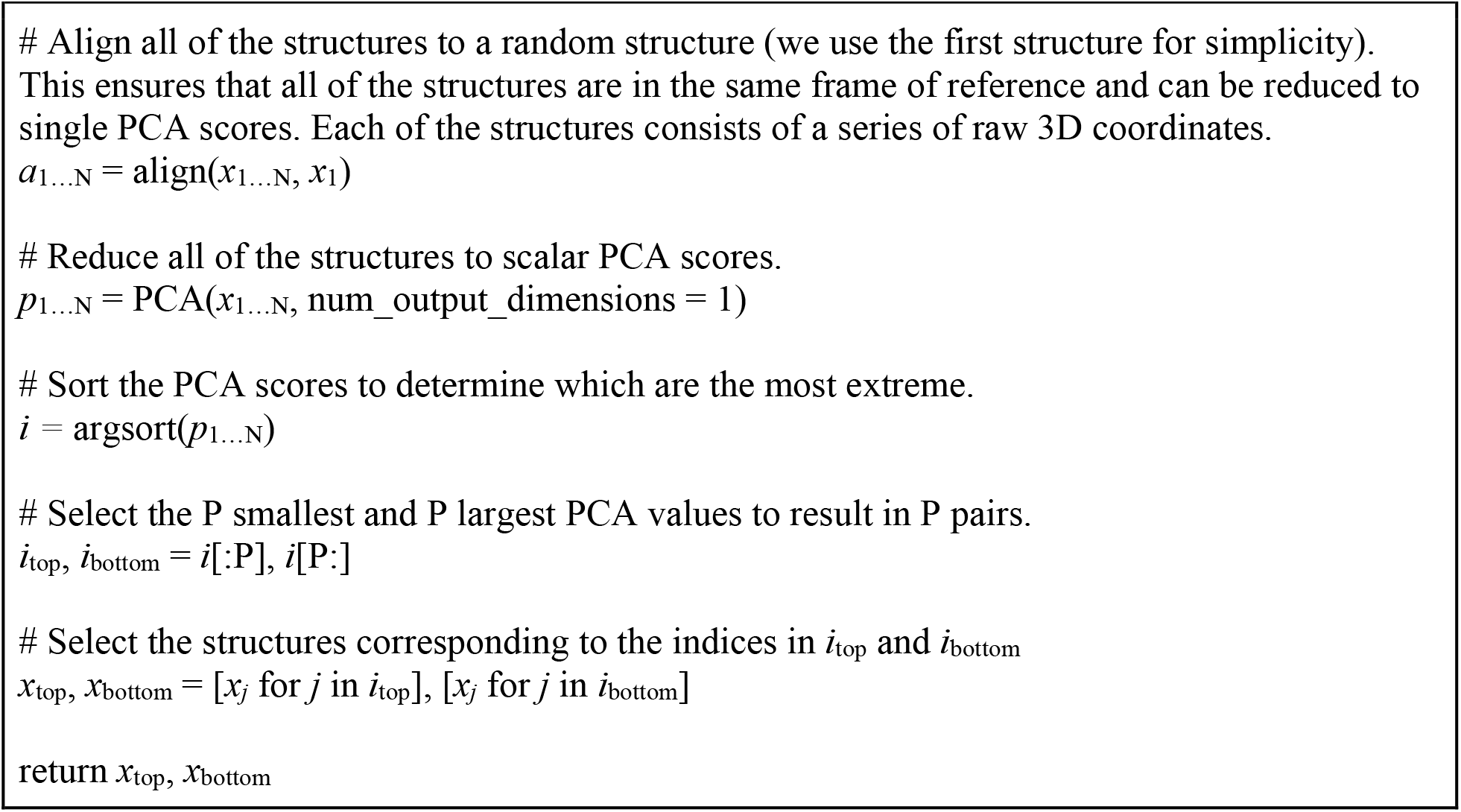

### Procedure of distogram-guided prediction

We implemented our entropy guidance method on top of OpenFold (a reproduction of AlphaFold2 in PyTorch)^36^, following the procedure outlined in Algorithm 2. The procedure involves a back propagation step that modifies the intermediate *m* and *z* representations using a custom implemented entropy loss before generating a final predicted structure. Our implementation uses gradient checkpointing and bfloat16 precision in order to reduce GPU memory requirements. For each target protein, we randomly selected one sequence from the pair of structures as the input sequence for prediction. We first ran AlphaFold2 without entropy guidance, followed by one step of entropy guided prediction using the Adam optimizer with a learning rate 0.01. No recycling was used. Both predicted structures before and after entropy guidance were taken as final output predictions. Ten prediction runs of AlphaFold2 were carried out using the originally available AlphaFold2 model presets released by DeepMind (model_{1-5} and model_{1-5}_ptm), resulting in a total of 20 structures per target protein (with 2 structures per run). The predictions were performed on 1-2 NVIDIA V100 GPUs.

We first evaluated our method on a set of 24 membrane proteins, each with experimentally determined structures in more than one conformation. Since AlphaFold2’s training cutoff date was 2018-04-30, structures released in the PDB after this date are not in its training dataset. These 24 proteins were categorized into two groups: one consisting of 12 proteins, with both conformations present in the training dataset, and the other consisting of 12 proteins with at least one conformation after the training cutoff date. Furthermore, we curated an independent test set of 13 membrane proteins, all of which were released in the PDB after the AlphaFold2 training date cutoff.

For AF-Cluster, we used the publicly available code at https://github.com/HWaymentSteele/AF_Cluster to cluster the MSA, and ran with default settings, using AF2 model_3_ptm in the OpenFold codebase^36^.

For a pair of the ground truth experimental structures GT_1_ and GT_2_, representing two conformations, and predicted structures, P_1…N_, a prediction is considered successful when the following two conditions are met:

1. There exists a P_i_ such that RMSD(P_i_, GT_1_) < 2.0 Å and RMSD(P_i_, GT_1_) < RMSD(P_i_, GT_2_)
2. There exists a P_i_ such that RMSD(P_i_, GT_2_) < 2.0 Å and RMSD(P_i_, GT_2_) < RMSD(P_i_, GT_1_)

The best prediction for each ground truth conformation is selected as the one that meets the above conditions and has the lowest RMSD to the corresponding ground truth conformation.

To identify potential conformational states from available experimental structures for a target protein, we searched the PDB for sequences with a sequence identity threshold of 90% to the original sequence, limiting search results to 100 per query. We then clustered the returned PDBs using disjoint set union with a distance threshold of 1.0 Å (i.e. if two protein structures are within 1.0 Å, then they are assigned to the same cluster). For each method (AF-Cluster, AF2, EGF), we used the following criteria to identify pairs of PDBs for each target protein:

1. For each cluster, we determined the minimum RMSD between any cluster member and the predicted structures.
2. We then selected two PDBs from the cluster representatives that are more than 2.0 Å apart and that are within 2.0 Å of a prediction.
3. For certain target proteins with conformations that are close to each other, we manually selected the pair of PDBs and examined the experimental structures to exclude disordered/irrelevant N-/C-terminal regions or disordered loops. Details on the specific conformations and sequences used are provided below in section “Details on Sequences Used”.

For target proteins where a method only predicts fewer than two conformations, we still provide two conformations for informational purposes. Detailed lists for each method on each dataset are given below:

AF-Cluster on 24 protein dataset:

| Sequence | PDB #1 | PDB #2 |
| --- | --- | --- |
| 5BZ2_A | 4BWZ_A | 5BZ2_A |
| 1XFH_A | 6X16_C | 6UWL_A |
| 4QIQ_A | 4GBY_A | 4JA4_C |
| 6LYY_A | 7CKO_A | 7CKR_A |
| 7LIA_A | 7TXT_S | 6DZZ_A |
| 3B4R_B | 3B4R_A | 3B4R_B |
| 2NRF_A | 6PJU_A | 2NRF_A |
| 8DKI_P | 8DKW_P | 8DYP_A |
| 4ZP0_A | 6EUQ_A | 6GV1_A |
| 4N7X_A | 7CYK_B | 4N7X_A |
| 6GCT_A | 8OUJ_C | 8OUH_B |
| 2X79_A | 4D1C_A | 2X79_A |
| 3TT1_A | 7DJC_A | 6XWM_B |
| 6GCI_A | 4C9G_A | 6GCI_A |
| 7WMV_A | 7WMV_A | 7SLA_A |
| 5AY0_A | 5AYM_A | 5AY0_A |
| 5L2A_C | 5L26_B | 5U9W_C |
| 1PV7_A | 2Y5Y_A | 6VBG_A |
| 6FHZ_A | 3WBN_A | 6FHZ_A |
| 7AAR_A | 6H7D_A | 7AAR_A |
| 8EX4_A | 8EX6_A | 8EX7_A |
| 6NC9_A | 6NC6_B | 6NC8_A |
| 4YBQ_A | 4YBQ_A | 4YB9_D |
| 7QPA_A | 7QPC_B | 7QPA_B |

AF2 baseline on 24 protein dataset:

| Sequence | PDB #1 | PDB #2 |
| --- | --- | --- |
| 5BZ2_A | 4BWZ_A | 5BZ2_A |
| 1XFH_A | 6X16_C | 6V8G_C |
| 4QIQ_A | 4JA4_C | 4GBY_A |
| 6LYY_A | 7CK0_A | 6LYY_A |
| 7LIA_A | 7TXT_S | 6DZZ_A |
| 3B4R_B | 3B4R_A | 3B4R_B |
| 2NRF_A | 6PJU_A | 2NRF_A |
| 8DKI_P | 8DYP_A | 8DKX_P |
| 4ZP0_A | 6EUQ_A | 6GV1_A |
| 4N7X_A | 7CYK_B | 6LH1_A |
| 6GCT_A | 8OUJ_B | 8OUJ_C |
| 2X79_A | 4D1C_A | 2X79_A |
| 3TT1_A | 7DIX_A | 6XWM_A |
| 6GCI_A | 4C9G_A | 6GCI_A |
| 7WMV_A | 7WMV_A | 7SLA_A |
| 5AY0_A | 5AYM_A | 6BTX_A |
| 5L2A_C | 5L2B_A | 5L27_C |
| 1PV7_A | 2Y5Y_A | 4ZYR_B |
| 6FHZ_A | 3VVN_A | 6FHZ_A |
| 7AAR_A | 7AAR_A | 6H7D_A |
| 8EX4_A | 8EX6_A | 8EX8_A |
| 6NC9_A | 5T77_A | 6NC7_B |
| 4YBQ_A | 4YB9_D | 4YBQ_A |
| 7QPA_A | 7QPC_B | 7QP9_B |

EGF on 24 protein dataset:

| Sequence | PDB #1 | PDB #2 |
| --- | --- | --- |
| 5BZ2_A | 4BWZ_A | 5BZ2_A |
| 1XFH_A | 6X16_C | 6BMI_B |
| 4QIQ_A | 4JA4_C | 6N3I_A |
| 6LYY_A | 7CK0_A | 7CKR_A |
| 7LIA_A | 7TXT_S | 6DZZ_A |
| 3B4R_B | 3B4R_A | 3B4R_B |
| 2NRF_A | 6PJU_A | 2NRF_A |
| 8DKI_P | 8DYP_A | 8DKX_P |
| 4ZP0_A | 6EUQ_A | 6GV1_A |
| 4N7X_A | 7CYK_B | 6LH1_A |
| 6GCT_A | 8OUJ_B | 8OUJ_C |
| 2X79_A | 4D1C_A | 2X79_A |
| 3TT1_A | 7DIX_A | 6XWM_A |
| 6GCI_A | 4C9G_A | 6GCI_A |
| 7WMV_A | 7WMV_A | 7SLA_A |
| 5AY0_A | 5AYM_A | 6BTX_A |
| 5L2A_C | 5L2B_A | 5L2B_C |
| 1PV7_A | 5GXB_A | 2Y5Y_A |
| 6FHZ_A | 3VVN_A | 6FHZ_A |
| 7AAR_A | 7AAR_A | 6H7D_A |
| 8EX4_A | 8EX6_A | 8EX7_A |
| 6NC9_A | 5T77_A | 6NC9_A |
| 4YBQ_A | 4YB9_D | 4YBQ_A |
| 7QPA_A | 7QPC_B | 7QP9_B |

AF2 baseline on 13 protein dataset:

| Sequence | PDB #1 | PDB #2 |
| --- | --- | --- |
| 8VZN_A | 8VZN_A | 8VZO_A |
| 8WNT_B | 8WNY_B | 8WNT_B |
| 8J76_A | 8J74_A | 8J77_A |
| 8WFI_A | 8WFK_A | 6ZPL_A |
| 8WM7_A | 8WM7_A | 8W9M_A |
| 8JT9_A | 8JT5_A | 8X09_A |
| 8TGG_B | 8TGI_A | 8TGG_B |
| 7PQG_A | 7PQG_A | 7VAG_A |
| 8JZU_A | 8JZU_A | 8P6A_A |
| 8SC2_A | 8SC3_A | 8JTT_A |
| 8I3C_A | 8WBX_B | 8WAM_B |
| 8J7Y_B | 8J7X_B | 8J7W_B |
| 8W6T_A | 8W6T_A | 8W6T_B |

EGF on 13 protein dataset:

| Sequence | PDB #1 | PDB #2 |
| --- | --- | --- |
| 8VZN_A | 8VZN_A | 8VZO_A |
| 8WNT_B | 8WNY_B | 8WNT_B |
| 8J76_A | 8J74_A | 8J76_A |
| 8WFI_A | 8WFK_A | 6ZPL_A |
| 8WM7_A | 8WM7_A | 8W9M_A |
| 8JT9_A | 8JT5_A | 8X09_A |
| 8TGG_B | 8TGI_A | 8TGG_B |
| 7PQG_A | 7PQG_A | 7VAG_A |
| 8JZU_A | 8JZU_A | 8P6A_A |
| 8SC2_A | 8SC3_A | 8JTZ_A |
| 8I3C_A | 8WBX_B | 8I3B_B |
| 8J7Y_B | 8J7X_B | 8J7W_B |
| 8W6T_A | 8W6T_A | 8W6T_B |

### DBSCAN clustering

For DBSCAN clustering, we used a precomputed distance matrix with pairwise RMSDs between predicted structures, with epsilon of 1.0 Å and min_samples of 1, using the implementation from scikit-learn, available at https://scikit-learn.org/dev/modules/generated/sklearn.cluster.DBSCAN.html

### RMSD calculation

To effectively align and calculate RMSD between multiple pairs, we adapted the sequence alignment script from the Molstar viewer^37^ (https://github.com/molstar/molstar) to Cython^38^ and integrated it with a Kabsch RMSD^32-34^ calculator from https://github.com/Bernhard10/py_qcprot (Supplementary information). Our custom RMSD method also enables preloading multiple structures and calculating pairwise RMSD between many pairs with high efficiency.

### Details on Sequences Used

#### Dataset with 24 proteins

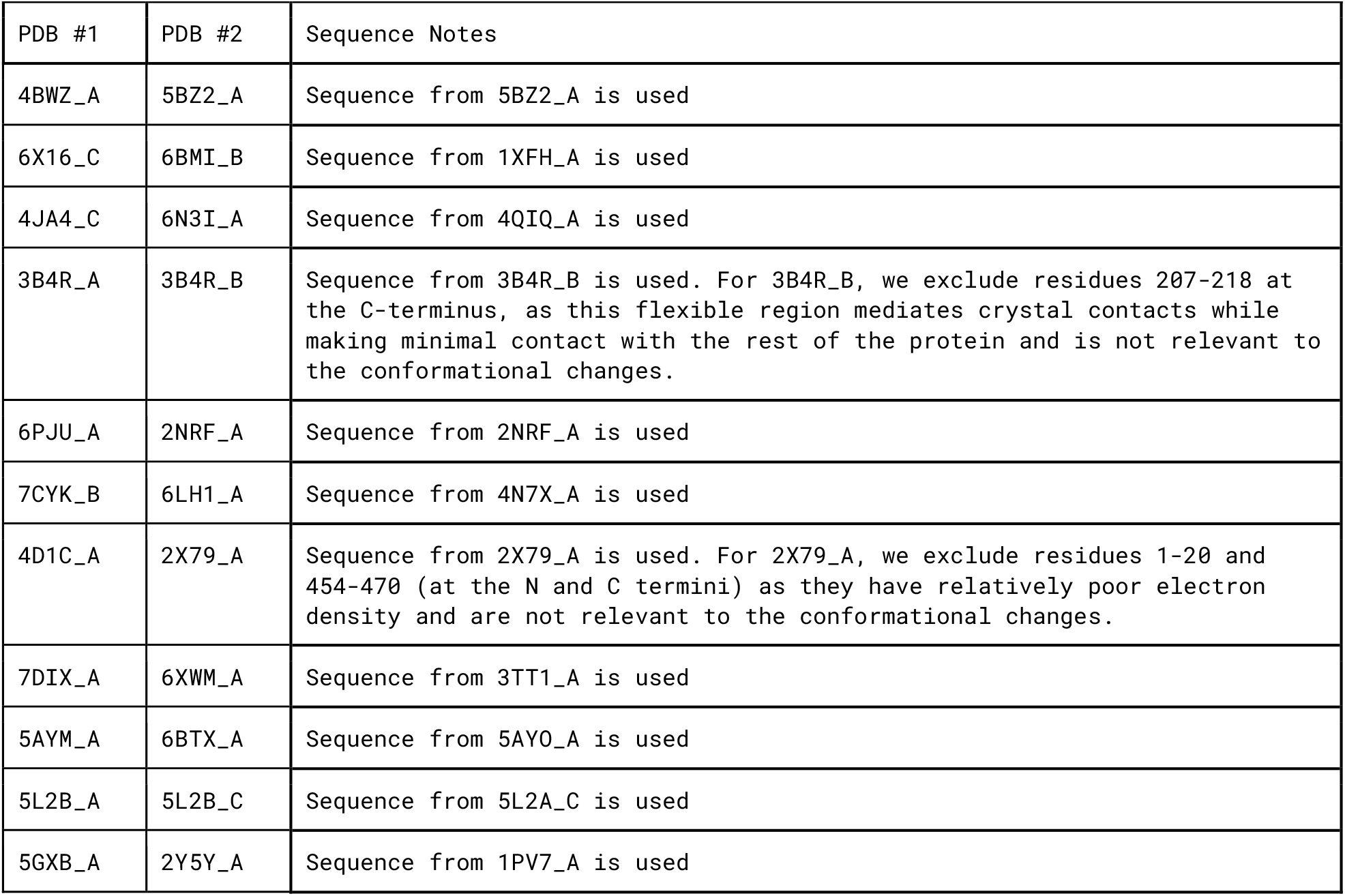
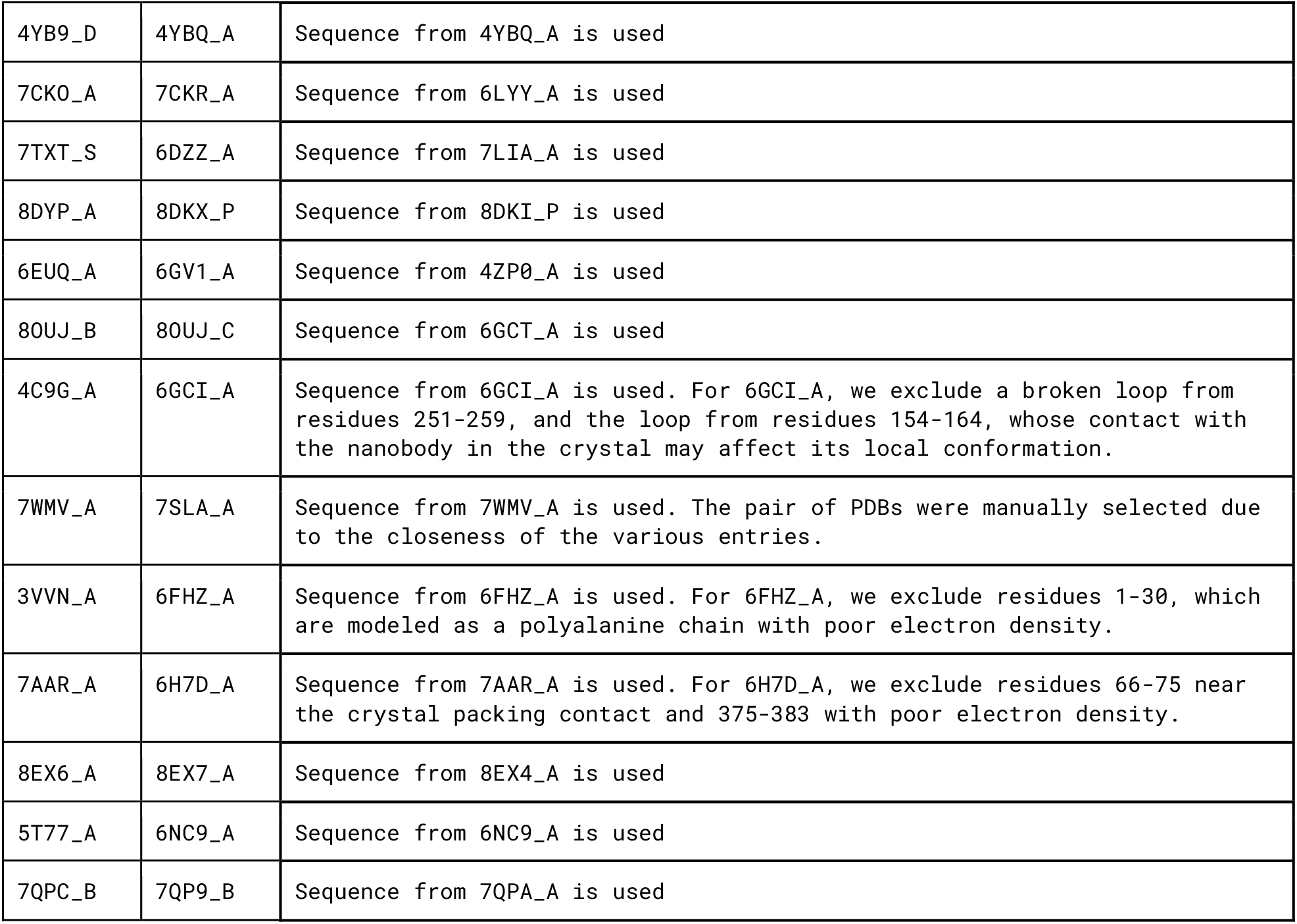

### Dataset with 13 proteins

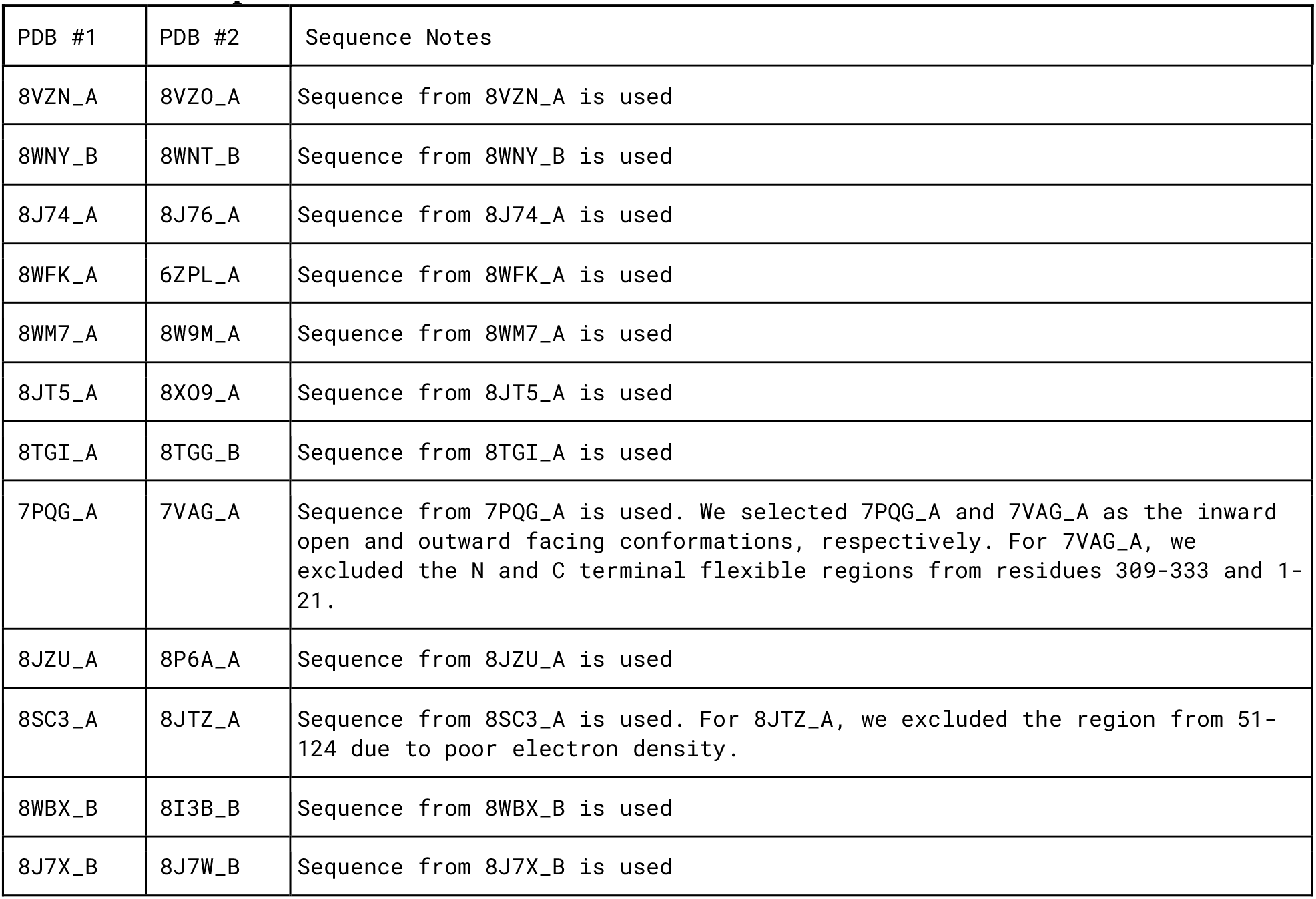
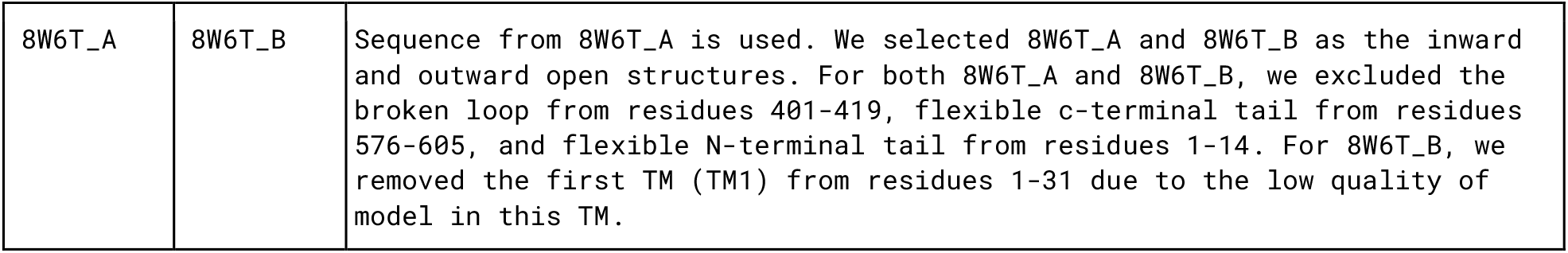

| PDB #1 | PDB #2 | Sequence Notes |
| --- | --- | --- |
| 8VZN_A | 8VZ0_A | Sequence from 8VZN_A is used |
| 8WNY_B | 8WNT_B | Sequence from 8WNY_B is used |
| 8J74_A | 8J76_A | Sequence from 8J74_A is used |
| 8WFK_A | 6ZPL_A | Sequence from 8WFK_A is used |
| 8WM7_A | 8W9M_A | Sequence from 8WM7_A is used |
| 8JT5_A | 8X09_A | Sequence from 8JT5_A is used |
| 8TGI_A | 8TGG_B | Sequence from 8TGI_A is used |
| 7PQG_A | 7VAG_A | Sequence from 7PQG_A is used. We selected 7PQG_A and 7VAG_A as the inward open and outward facing conformations, respectively. For 7VAG_A, we excluded the N and C terminal flexible regions from residues 309-333 and 1-21. |
| 8JZU_A | 8P6A_A | Sequence from 8JZU_A is used |
| 8SC3_A | 8JTZ_A | Sequence from 8SC3_A is used. For 8JTZ_A, we excluded the region from 51-124 due to poor electron density. |
| 8WBX_B | 8I3B_B | Sequence from 8WBX_B is used |
| 8J7X_B | 8J7W_B | Sequence from 8J7X_B is used |
| 8W6T_A | 8W6T_B | Sequence from 8W6T_A is used. We selected 8W6T_A and 8W6T_B as the inward and outward open structures. For both 8W6T_A and 8W6T_B, we excluded the broken loop from residues 401-419, flexible c-terminal tail from residues 576-605, and flexible N-terminal tail from residues 1-14. For 8W6T_B, we removed the first TM (TM1) from residues 1-31 due to the low quality of model in this TM. |

## Code availability

The code for EGF is available at: www.github.com/Fenglaboratory/EGF

## Acknowledgements

We thank J. Zhang and B. Rao for helpful discussions, and A. Brunger for helpful suggestions. L.F. acknowledges the support from Stanford University and NIH-R35GM153424.

## Author Contributions

D.W. and L.F. conceived the project. D.W. carried out research. D.W. and L.F. wrote the manuscript. L.F. supervised the project.

## Additional information

