## Supplementary information for "Robust Prediction of Multiple Protein Conformations with Entropy Guidance"

The sequence alignment code was adapted from Molstar viewer[32] (https://github.com/molstar/molstar) into Cython. It was then combined with a RMSD calculator using Kabsch algorium[33-35]. (https://github.com/Bernhard10/py_qcprot).

The alignment code is given below:

Code Sample 1: Sequence Alignment in Cython

```
# distutils: language=c++
#cython: language_level=3
from libcpp.vector cimport vector

cdef int INF = 1000000000
cdef int[2000][2000] S
cdef int[2000][2000] V
cdef int[2000][2000] H
cdef int gapPenalty = -11, gapExtensionPenalty = -1

blosum62 = [
   # A  R  N  D  C  Q  E  G  H  I  L  K  M  F  P  S  T  W  Y  V  B  Z  X
   [4, -1, -2, -2, 0, -1, -1, 0, -2, -1, -1, -1, -1, -2, -1, 1, 0, -3, -2, 0, -2, -1, 0], # A
   [-1, 5, 0, -2, -3, 1, 0, -2, 0, -3, -2, 2, -1, -3, -2, -1, -1, -3, -2, -3, -1, 0, -1], # R
   [-2, 0, 6, 1, -3, 0, 0, 0, 1, -3, -3, 0, -2, -3, -2, 1, 0, -4, -2, -3, 3, 0, -1], # N
   [-2, -2, 1, 6, -3, 0, 2, -1, -1, -3, -4, -1, -3, -3, -1, 0, -1, -4, -3, -3, 4, 1, -1], # D
   [0, -3, -3, -3, 9, -3, -4, -3, -3, -1, -1, -3, -1, -2, -3, -1, -1, -2, -2, -1, -3, -3, -2], # C
   [-1, 1, 0, 0, -3, 5, 2, -2, 0, -3, -2, 1, 0, -3, -1, 0, -1, -2, -1, -2, 0, 3, -1], # Q
   [-1, 0, 0, 2, -4, 2, 5, -2, 0, -3, -3, 1, -2, -3, -1, 0, -1, -3, -2, -2, 1, 4, -1], # E
   [0, -2, 0, -1, -3, -2, -2, 6, -2, -4, -4, -2, -3, -3, -2, 0, -2, -2, -3, -3, -1, -2, -1], # G
   [-2, 0, 1, -1, -3, 0, 0, -2, 8, -3, -3, -1, -2, -1, -2, -1, -2, -2, 2, -3, 0, 0, -1], # H
   [-1, -3, -3, -3, -1, -3, -3, -4, -3, 4, 2, -3, 1, 0, -3, -2, -1, -3, -1, 3, -3, -3, -1], # I
   [-1, -2, -3, -4, -1, -2, -3, -4, -3, 2, 4, -2, 2, 0, -3, -2, -1, -2, -1, 1, -4, -3, -1], # L
   [-1, 2, 0, -1, -3, 1, 1, -2, -1, -3, -2, 5, -1, -3, -1, 0, -1, -3, -2, -2, 0, 1, -1], # K
   [-1, -1, -2, -3, -1, 0, -2, -3, -2, 1, 2, -1, 5, 0, -2, -1, -1, -1, -1, 1, -3, -1, -1], # M
   [-2, -3, -3, -3, -2, -3, -3, -3, -1, 0, 0, -3, 0, 6, -4, -2, -2, 1, 3, -1, -3, -3, -1], # F
   [-1, -2, -2, -1, -3, -1, -1, -2, -2, -3, -3, -1, -2, -4, 7, -1, -1, -4, -3, -2, -2, -1, -2], # P
   [1, -1, 1, 0, -1, 0, 0, 0, -1, -2, -2, 0, -1, -2, -1, 4, 1, -3, -2, -2, 0, 0, 0], # S
   [0, -1, 0, -1, -1, -1, -1, -2, -2, -1, -1, -1, -1, -2, -1, 1, 5, -2, -2, 0, -1, -1, 0], # T
   [-3, -3, -4, -4, -2, -2, -3, -2, -2, -3, -2, -3, -1, 1, -4, -3, -2, 11, 2, -3, -4, -3, -2], # W
   [-2, -2, -2, -3, -2, -1, -2, -3, 2, -1, -1, -2, -1, 3, -3, -2, -2, 2, 7, -1, -3, -2, -1], # Y
   [0, -3, -3, -3, -1, -2, -2, -3, -3, 3, 1, -2, 1, -1, -2, -2, 0, -3, -1, 4, -3, -2, -1], # V
   [-2, -1, 3, 4, -3, 0, 1, -1, 0, -3, -4, 0, -3, -3, -2, 0, -1, -4, -3, -3, 4, 1, -1], # B
   [-1, 0, 0, 1, -3, 3, 4, -2, 0, -3, -3, 1, -1, -3, -1, 0, -1, -3, -2, -2, 1, 4, -1], # Z
   [0, -1, -1, -1, -2, -1, -1, -1, -1, -1, -1, -1, -1, -1, -2, 0, 0, -2, -1, -1, -1, -1, -1] # X
]

aminoacids = 'ARNDCQEGHILKMFPSTWYVBZX'

substitution_mapping = {
   amino1: {
       amino2: blosum62[i][j] for j, amino2 in enumerate(aminoacids)
   } for i, amino1 in enumerate(aminoacids)
}

def align(seqA: str, seqB: str, return_visual: bool = False):
   n = len(seqA)
   m = len(seqB)

   for i in range(n + 1):
       S[i][0] = gapPenalty
       H[i][0] = -INF

   for j in range(m + 1):
       S[0][j] = gapPenalty
       V[0][j] = -INF

   S[0][0] = 0
```

```
score_fn = lambda i, j: substitution_mapping[seqA[i]][seqB[j]] # 5 if seqA[i] == seqB[j] else -3

for i in range(1, n + 1):
    for j in range(1, m + 1):
        V[i][j] = max(
            S[i - 1][j] + gapPenalty,
            V[i - 1][j] + gapExtensionPenalty
        )

        H[i][j] = max(
            S[i][j - 1] + gapPenalty,
            H[i][j - 1] + gapExtensionPenalty
        )

        S[i][j] = max(
            S[i - 1][j - 1] + score_fn(i - 1, j - 1),
            V[i][j],
            H[i][j]
        )

i = n
j = m

aliA = []
aliB = []

if S[i][j] >= V[i][j]:
    mat = 'S'
    score = S[i][j]
elif V[i][j] >= H[i][j]:
    mat = 'V'
    score = V[i][j]
else:
    mat = 'H'
    score = H[i][j]

indA, indB = [], []
while i > 0 and j > 0:
    if mat == 'S':
        if S[i][j] == S[i - 1][j - 1] + score_fn(i - 1, j - 1):
            aliA.append(seqA[i - 1])
            aliB.append(seqB[j - 1])
            indA.append(i-1)
            indB.append(j-1)
            i -= 1
            j -= 1
            mat = 'S'
        elif S[i][j] == V[i][j]:
            mat = 'V'
        elif S[i][j] == H[i][j]:
            mat = 'H'
        else:
            i -= 1
            j -= 1
    elif mat == 'V':
        if V[i][j] == V[i - 1][j] + gapExtensionPenalty:
            aliA.append(seqA[i - 1])
            aliB.append('-')
            i -= 1
            mat = 'V'
        elif V[i][j] == S[i - 1][j] + gapPenalty:
            aliA.append(seqA[i - 1])
            aliB.append('-')
            i -= 1
            mat = 'S'
        else:
            i -= 1
    elif mat == 'H':
        if H[i][j] == H[i][j - 1] + gapExtensionPenalty:
            aliA.append('-')
            aliB.append(seqB[j - 1])
```

```
            j -= 1
            mat = 'H'
        elif H[i][j] == S[i][j - 1] + gapPenalty:
            aliA.append('-')
            aliB.append(seqB[j - 1])
            j -= 1
            mat = 'S'
        else:
            j -= 1

    while i > 0:
        aliA.append(seqA[i - 1])
        aliB.append('-')
        i -= 1

    while j > 0:
        aliA.append('-')
        aliB.append(seqB[j - 1])
        j -= 1

    aliA = ''.join(aliA[::-1])
    aliB = ''.join(aliB[::-1])

    if return_visual:
        return aliA, aliB
    else:
        return indA[::-1], indB[::-1]
```
